# Functional redundancy and formin-independent localization of tropomyosin isoforms in Saccharomyces cerevisiae

**DOI:** 10.1101/2024.04.04.587703

**Authors:** Anubhav Dhar, VT Bagyashree, Sudipta Biswas, Jayanti Kumari, Amruta Sridhara, B Jeevan Subodh, Shashank Shekhar, Saravanan Palani

## Abstract

Tropomyosin is an actin binding protein which protects actin filaments from cofilin-mediated disassembly. Distinct tropomyosin isoforms have long been hypothesized to differentially sort to subcellular actin networks and impart distinct functionalities. Nevertheless, a mechanistic understanding of the interplay between Tpm isoforms and their functional contributions to actin dynamics has been lacking. In this study, we present and charcaterize mNeonGreen-Tpm fusion proteins that exhibit good functionality in cells as a sole copy, surpassing limitations of existing probes and enabling real-time dynamic tracking of Tpm-actin filaments *in vivo*. Using these functional Tpm fusion proteins, we find that *S. cerevisiae* Tpm isoforms, Tpm1 and Tpm2, colocalize on actin cables and indiscriminately bind to actin filaments nucleated by either formin isoform-Bnr1 and Bni1 *in vivo*, in contrast to the long-held paradigm of Tpm-formin pairing. We show that cellular Tpm levels regulate endocytosis by affecting balance between linear and branched actin networks in yeast cells. Finally, we discover that Tpm2 can protect and organize functional actin cables in absence of Tpm1. Overall, our work supports a concentration-dependent and formin isoform independent model of Tpm isoform binding to F-actin and demonstrates for the first time, the functional redundancy of the paralog Tpm2 in actin cable maintenance in *S. cerevisiae*.

## Introduction

Tropomyosins (Tpm) are a major class of actin-binding proteins present in fungi^1^ and metazoans^2–4^. Tropomyosins are helical coiled-coil proteins that form head-to-tail dimers and co-polymerize with actin filaments^5–8^. Tpm was initially discovered as a component of muscle^9^ where it regulates actomyosin contraction^10,11^. Later studies revealed that Tpm isoforms are also abundant in non-muscle cells^4,12–14^ and even single-celled eukaryotic organisms like fungi^13,15,16^. The canonical role of non-muscle Tpm is to protect actin filaments from actin-severing proteins like cofilin^17–19^ and thus regulate turnover of actin networks^20^. Tpms display a great diversity of isoforms across species with involvement in various cellular processes such as cell motility^21^, cell division^13^, organellar movement, secretory vesicle delivery, etc.^2–4,22^. Mammals have four tropomyosin genes which produce around 40 isoforms via alternate splicing^2,14,23^. Various studies have shown that Tpm isoforms sort extensively to distinct actin filament subpopulations and define diverse functionalities for these networks by controlling the interactions between F-actin and other actin-binding proteins such as myosin, cofilin, capping protein, etc.^2,4,24–29^. The molecular basis for Tpm isoform sorting remains an unresolved question with various mechanisms such as concentration dependence^30^ or a formin-dependent targeting^31,32^ having been observed. Structural analysis using cryoEM has now revealed that Tpm isoforms may exert differential functions via different modes of binding and controlling accessibility of ABPs like myosin and cofilin to F-actin^33,34^. How Tpm isoforms have been fine-tuned to perform distinct spatial sorting and functions during evolution remains an active area of research and their crosstalk during recruitment and maturation within the same actin structure is just beginning to be understood^21,35^.

The eukaryotic model organism *Saccharomyces cerevisiae* has two tropomyosin isoforms encoded by two different genes-*tpm1* and *tpm2*^1,15,16^. Tpm1 (199 amino acids) is considered a duplicated paralog of Tpm2 (161 amino acids) with a 38 amino acid internal duplication^16^. Tpm1 is considered the major isoform and Δ*tpm1* cells display a near complete loss of actin cables and lethality at higher temperatures^13,28,29^. Δ*tpm2* cells on the other hand show no detectable changes in actin cable organization and cell growth^16,36^. Interestingly, deletion of both *tpm1* and *tpm2* is synthetic lethal^16^. Cells compromised in actin severing factors such as Δ*aip1* or Δ*srv2* mutants show partial restoration of cables in Δ*tpm1* cells which suggests that Tpm1 is majorly involved in protection of the actin filaments from severing proteins of the CCA (Cofilin-Coronin-Aip1) complex *in vivo*^37,38^. The minor isoform Tpm2 is expressed at ∼5 fold lower levels and Tpm2 overexpression could not restore normal growth and actin cables in Δ*tpm1* cells in a previous study^16^. The *in vivo* functions of Tpm2 have remained unclear but studies have suggested a role for Tpm2 in negatively controlling retrograde actin cable flow (RACF) possibly via regulating actin filament binding of the type-II myosin Myo1, which is positive regulator of RACF^39,40^. Both Tpm1 and Tpm2 facilitate processive motion of the type-V myosin Myo2 on coated actin filaments *in vitro*^41^, contributing to the anterograde flow of organelles and cargo to the growing bud^22,42,43^. Thus, Tpm1 is believed to play a major role in maintaining the actin cytoskeleton in yeast while the functions of Tpm2 are believed to be majorly distinct from Tpm1^16^. While genetic studies have indicated a possible preference between Tpm isoforms and formin isoforms^36^, whether Tpm1 and Tpm2 localize to distinct actin filaments/cables in *S. cerevisiae* has remained unaddressed mainly due to absence of functional Tpm fluorescent fusions and live-cell imaging data for Tpm1 and Tpm2. Recent work from our lab made it possible to visualize the live dynamics of Tpm across species using mNG-Tpm fusion proteins^44^ for the first time and revealed their localization to actin cables and the actomyosin ring during the cell cycle^44^. However, their combined action and roles in actin cable stability *in vivo* is not understood.

In this study, we assess the functionality of mNeonGreen-Tpm fusion proteins and show that mNG-Tpm fusion proteins are functional and restore viability as sole copies in cells lacking native Tpm1 and Tpm2. We also engineer -AS-dipeptide containing mNG-^AS^Tpm constructs to account for lack of N-terminal acetylation and improve functionality *in vivo*. Using mNG-^AS^Tpm fusion proteins, we find that Tpm1 and Tpm2 indiscriminately associate with actin filaments made by either formin Bnr1 and Bni1, unlike the differential association of acetylated and unacetylated Cdc8 to distinct formin-nucleated filament populations shown in *S. pombe*^31,45^. We report a novel function of Tpm2 and show that Tpm2 can compensate for loss of Tpm1 upon increased expression *in vivo.* In contrast to the long-held view, we find that increased Tpm2 expression can restore full length actin cables and actin-cable dependent functions such as vesicle targeting to bud and maintenance of normal mitochondrial morphology suggesting that Tpm2 can independently organize a functional actin cable network in *S. cerevisiae* in the absence of Tpm1. Lastly, we also report the role of Tpm1 and Tpm2 in maintaining linear-to-branched actin network homeostasis in yeast, revealing an intriguing cascading effect of Tpm on actin dynamics at sites of endocytosis. Overall, our findings support a concentration-dependent and formin-independent localization of Tpm isoforms to actin cables and unveil the functional redundancy between Tpm isoforms in *S. cerevisiae*.

## Results

### mNG-Tpm fusion proteins are functionally-active tagged Tropomyosin fusion proteins

Visualization of tropomyosin in live cells has been a major challenge because fluorescently-tagged tropomyosin fusions are not completely functional and thus, need careful interpretation for accurate insights about Tpm localization and dynamics *in vivo*^46^. We have recently shown that mNeonGreen-Tpm (mNG-Tpm) fusion proteins clearly report localization of Tpm isoforms to actin structures *in vivo* in fungal and mammalian cells^44^ **(Fig. S1A, S1B)** but their functionality compared to native Tpm remained unknown. To address if mNG-Tpm fusion proteins could compensate for loss of native Tpm in cells, we assessed complementation of cellular defects caused by deletion of *tpm1* gene by mNG-Tpm1 expressed under the native *tpm1* promoter. We found that mNG-Tpm1 expression could near completely restore normal growth **(Fig. S1C, S1D)** but only partially restore the length of actin cables in Δ*tpm1* cells which exhibit slower growth and absence of actin cables **(Fig. S1G, 1C)**. The presence of shorter actin cables suggests partial functionality of mNG-Tpm1 fusion as compared to the native Tpm1 protein and that partial length cables are sufficient to majorly support normal growth via proper cargo targeting to the bud. One possible explanation for presence of shorter actin cables is the lack of N-terminal acetylation of mNG-Tpm by the Nat3-Mdm20 complex which is essential for normal Tpm-actin binding *in vivo*^47–49^. This could result in reduced binding affinities towards F-actin as compared to native Tpm. Thus, we constructed mNG-^AS^Tpm, a modified mNG-Tpm fusion protein with an Ala-Ser dipeptide before the starting methionine of Tpm that has been routinely used to restore normal binding affinities by stabilizing the N-terminal alpha-helix in the absence of the N-terminal acetylation^25,36,50–52^. Live-cell imaging of cells expressing mNG-^AS^Tpm fusion protein as an integrated single copy showed improved but clear signal enhancement of both mNG-^AS^Tpm1 and mNG-^AS^Tpm2 on actin cables and the actomyosin ring (**Fig. 1A**, **S1H, Supplementary Movie 1-2)**. mNG-^AS^Tpm1 expression could completely rescue the growth defects **(Fig. S1C, S1D)** and restore full-length actin cables in Δ*tpm1* cells when expressed at similar levels as mNG-Tpm1 **(Fig. 1B, 1C)**. These results clearly indicate that mNG-^AS^Tpm1 is a functional tagged tropomyosin with similar activity as native Tpm1. To test for functionality of tagged Tpm2, we expressed mNG-Tpm1, mNG-^AS^Tpm1, mNG-Tpm2, mNG-^AS^Tpm2 under their native promoters in Δ*tpm1*Δ*tpm2* cells containing a copy of Tpm1 in a *ura3*-centromeric plasmid. Shuffling of these strains to 5’FOA-containing media revealed that all mNG-Tpm fusions rescued synthetic lethality of Δ*tpm1*Δ*tpm2* cells when expressed from a high-copy plasmid (*pRS425*) **(Fig. S1E)** but only mNG-Tpm1 and mNG-^AS^Tpm1 restored viability when expressed from single-copy integration plasmid (*pRS305*) **(Fig. S1F)**. This suggests that both mNG-Tpm1 and mNG-Tpm2 fusions are functional tagged Tpm fusions and Tpm1 tolerates the presence of the N-terminal tag better than Tpm2. Next, we tested whether actin cables restored by mNG-Tpm1 and mNG-^AS^Tpm1 are functional and assessed mitochondrial morphology as a readout of function. We found that mitochondrial fragment number is restored to wildtype levels by expression of both mNG-Tpm1 and mNG-^AS^Tpm1 in Δ*tpm1* cells which show hyper-fragmented mitochondria^53,54^, suggesting again that full length cables are not necessary for restoring downstream actin cable-dependent functions. These data also suggest that Tpm mediated actin cable stability *in vivo* may be one of the factors determining actin cable length control with cell size in yeast^55,56^ (**Fig. 1D**, **S1G)**.

**Figure 1:**
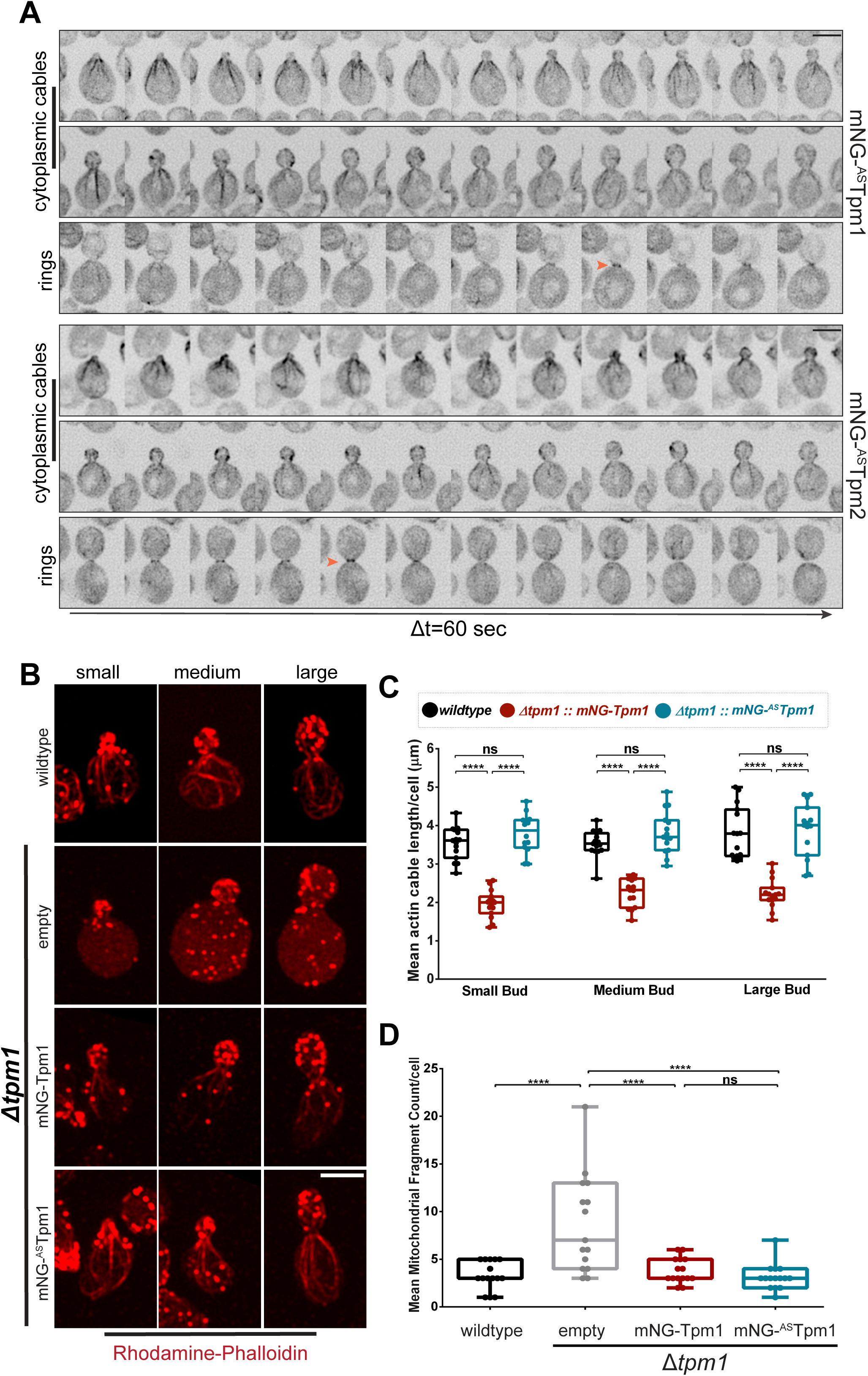
mNeonGreen-Tpm fusion proteins are functional and restore normal growth and ac-tin cytoskeleton organization *in vivo*. **(A)** Representative time-lapse montages of wildtype yeast cells expressing mNG-^AS^Tpm1 (top) or mNG-^AS^Tpm2 (bottom); scale bar -3μm. **(B)** Representative images of cells of indicated yeast strains stained with Rhodamine-phalloidin; scale bar -3μm. **(C)** Plot representing mean actin cable length per cell for indicated strains as per experiment shown in (B). **(D)** Plot representing mean mitochondrial fragment count per cell for indicated strains as per experi-ment shown in S1G. (Box represents 25th and 75th percentile, line represents median, whiskers represent minimum and maximum values; One-Way Anova with Tukey’s Multiple Comparisons test was used in (C) and (D), * p < 0.05, ** p < 0.01, *** p < 0.001, **** p < 0.0001)

Taken together, our results demonstrate that mNG-Tpm fusion proteins are functional, and acetylation-mimic improves their functionality in vivo, thus, expanding their applicability to probe a broad range of questions about Tpm isoform biology.

### Tpm1 and Tpm2 indiscriminately bind to actin cables nucleated by both formin isoforms

Tpm isoforms are known to sort to distinct actin filament structures in space and time in various organisms^4,4,24,32,32,33,45,57–61^ and the basis of this spatial sorting remains an enigmatic question^30,62^. N-terminal acetylated and unacetylated forms of the fission yeast Tpm, Cdc8, show a formin-dependent spatial sorting to distinct actin cable networks^31,45^. Contrasting results from studies in mammalian cells support both formin-mediated^31,32^ or relative concentration-dependent sorting mechanisms^30^. The presence of two Tpm isoforms maintained at distinct expression levels (Tpm1 expressed ∼5-6 fold higher than Tpm2)^16,63^ and previous observations showing that Δ*tpm1* exhibits distinct genetic interactions with either formin-Bnr1 and Bni1 indicates a possible crosstalk between Tpm and formin isoforms in *S. cerevisiae*^36^. Tpm1 also increased Bni1-mediated nucleation of actin filaments in an *in vitro* assay without having any effect on Bnr1-mediated nucleation in the same study^36^. However, whether these differences observed *in vitro* translate to measurable effects and possible formin-based spatial sorting of Tpm1 and Tpm2 *in vivo* remains unclear. To answer these questions, we investigated if Tpm1 and Tpm2 preferred actin cables nucleated by a specific formin isoform -Bnr1 or Bni1. Bnr1 and Bni1 localize to the bud neck and bud cortex respectively from G1 till onset of cytokinesis where they nucleate distinct sets of actin filaments^64–68^. Localization of both Tpm1 and Tpm2 on actin cables could be observed in both Δ*bnr1* and Δ*bni1* cells (**Fig. 2A**, **2D)**, qualitatively suggesting that both Tpm1 and Tpm2 bind to actin cables nucleated by either formin - Bnr1 and Bni1 indiscriminately.

**Figure 2:**
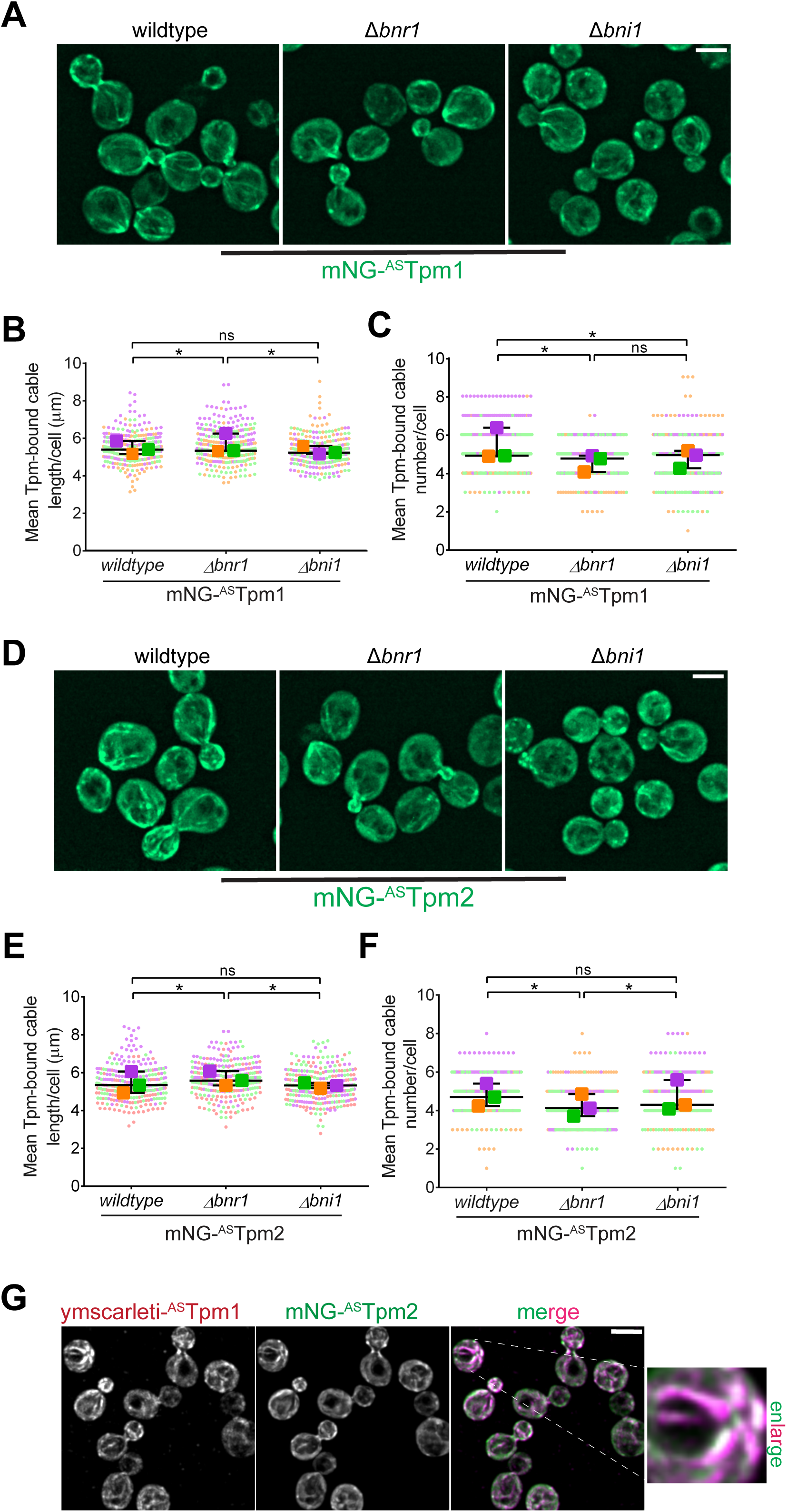
Tpm1 and Tpm2 colocalize and indiscriminately bind to actin filaments nucleated by formin isoforms, Bnr1 and Bni1. **(A)** Representative images of cells of wildtype, ∆*bnr1*, and ∆*bni1* cells expressing mNG-^AS^Tpm1 fusion protein; scale bar - 3μm. **(B)** Superplot representing mean Tpm-bound cable length per cell in wildtype, ∆*bnr1*, and ∆*bni1* cells expressing mNG-^AS^Tpm1; n=100 cells for each strain per replicate, N=3. **(C)** Superplot representing mean Tpm-bound cable number per cell in wildtype, ∆*bnr1*, and ∆*bni1* cells expressing mNG-^AS^Tpm1; n=100 cells for each strain per replicate, N=3. **(D)** Representative images of cells of wildtype, ∆*bnr1*, and ∆*bni1* cells expressing mNG-^AS^Tpm2 fusion protein; scale bar -3μm. **(E)** Superplot representing mean Tpm-bound cable length per cell in wildtype, ∆*bnr1*, and ∆*bni1* cells expressing mNG-^AS^Tpm2; n=100 cells for each strain per replicate, N=3. **(F)** Superplot representing mean Tpm-bound cable number per cell in wildtype, ∆*bnr1*, and ∆*bni1* cells expressing mNG-^AS^Tpm2; n=100 cells for each strain per replicate, N=3. **(G)** Representative images of wildtype yeast cells co-expressing ymScarleti-^AS^Tpm1 and mNG-^AS^Tpm2; scale bar - 3μm. (Superplots represent datapoints and means from three independent biological replicates marked in different colours; * p < 0.05, ** p < 0.01; One-Way Anova with Tukey’s Multiple Comparisons test was used in (B), (C), (E), (F); * p < 0.05, ** p < 0.01)

To assess for any minor biases in preference shown by Tpm1 and Tpm2 for formins Bnr1 and Bni1, we then performed an extensive quantitative analysis of actin cable organization visualized by phalloidin staining **(Fig. S2A, S2D)** and Tpm localization visualized by mNG-^AS^Tpm1/2 fusion proteins in wildtype, Δ*bnr1,* and Δ*bni1* cells expressed under their native promoters from an extra integrated copy at the *leu2* locus **(Fig. 2A, 2D)**. However, mNG-^AS^Tpm signal drastically weakened after fixation making it difficult to score properly for Tpm localization. Hence, we measured average actin and Tpm-bound actin cable length and number in these strains as a population average instead of simultaneous co-staining in single cells (**Fig. 2A, 2D, S2A, S2D**). Our analysis revealed that actin cable numbers dropped significantly upon deletion of either formin Bnr1 and Bni1 suggesting a redundant role for both formins in actin cable nucleation **(Fig. S2C, S2F),** while actin cable lengths were longer in *Δbnr1* as compared to both wildtype and Δ*bni1* cells **(Fig. S2B, S2E)** which could be explained by the elongated cell shape observed in Δ*bnr1* cells **(Fig. S2A, S2D)**. Both Tpm1-and Tpm2-bound cable length did not show a consistent signifiant change between wildtype, Δ*bnr1, and* Δ*bni1* cells (**Fig. 2B**, **2Ε)**. The average number of Tpm1-and Tpm2-bound cables per cell were reduced in Δ*bnr1 and* Δ*bni1* cells as compared to *wildtype* **(Fig. 2C, 2F)** but the reduction was comcomitant with the reduction in actin cable numbers observed in Δ*bnr1 and* Δ*bni1* **(Fig. S2C, S2F)** and no major dependence on either formin was observed for both Tpm1 and Tpm2. The ratio of mNG-^AS^Tpm1 and mNG-^AS^Tpm2 fluorescence in the bud to the mother compartment increased in Δ*bnr1* cells and decreased in Δ*bni1* cells as compared to wildtype cells **(Fig. S2G)**, consistent with the fact that Bnr1 nucelates cables only in the mother compartment and all cables present in the bud compartment are nucleated by Bni1 **(Fig. S2A, S2D)**. This suggests that Tpm1 and Tpm2 do not show preference for binding to cables in either bud or mother compartment as shown recently for another actin cable-binding protein, Abp140^69^. Since, mNG-^AS^Tpm1 is functional when expressed under native Tpm1 promoter **(Fig. S1D, S1E)**, we also conducted our analysis in strains lacking endogenous Tpm1 and expressing only mNG-^AS^Tpm1 to eliminate any effects of increased levels of functional Tpm1 protein. We found that mNG-^AS^Tpm1 still localized to actin cables in wildtype, Δ*bnr1, and* Δ*bni1* cells **(Fig. S3A)**. Tpm-bound actin cable length and Tpm-bound actin cable number was decreased in Δ*bnr1* and Δ*bni1* cells as compared to wildtype **(Fig. S3B, S3C)**, which was again concomitant with the decrease in actin cable length and number (**Fig. S3D, S3E, S3F**). Thus, our quantitative analysis also suggests that Tpm1 and Tpm2 binding to actin cables is unaffected by identity of the formin isoforms-Bnr1 and Bni1, implicating that they indiscriminately bind to both Bnr1-and Bni1-nucleated actin cables in *S. cerevisiae*.

Lastly, we assessed if Tpm1 and Tpm2 localize to the same set of actin cables *in vivo* by co-expressing ymScarleti-^AS^Tpm1 and mNG-^AS^Tpm2 fusion proteins. We observed that both ymScarleti-^AS^Tpm1 and mNG-^AS^Tpm2 localized to the same set of actin cables (**Fig. 2G**) in cells indicating that they may either co-polymerize on a single actin filament or are bound to different actin filaments bundled together into an actin cable.

Overall, our qualitative and quantitative analysis suggests that Tpm1 and Tpm2 indiscriminately bind to actin filaments nucleated by either formin, Bnr1 and Bni1, and do not exhibit spatial sorting to distinct actin cable networks in *S. cerevisiae*. Since, Tpm1 and Tpm2 have differential expression levels in cells, their relative local concentrations may be the major factor in determining their localization to actin filament networks in *S. cerevisiae* and governing their functions.

### Tpm1 and Tpm2 localize to the fusion focus in a Bnr1-independent manner in mating yeast cells

Polarized actin assembly is required for growth of the pheromone-induced mating projection (shmoo) in yeast cells^70–72^. Similar to polarized bud growth, formin-made actin cables act as tracks for vesicles to be delivered to the tip of the mating projection where coordinated exo-and endo-cytosis ensure polarized growth, membrane turnover, and cell wall remodelling^73,74^. It has been shown that Δ*tpm1* cells require much higher pheromone concentrations for shmoo formation as compared to wildtype cells^75^. To assess whether both Tpm1 and Tpm2 localized to the actin cables in the mating projection, we mated wildtype *MAT-a* haploid yeast cells expressing either mNG-^AS^Tpm1 or mNG-^AS^Tpm2 with wildtype *MAT-α* yeast cells. Both mNG-^AS^Tpm1 and mNG-^AS^Tpm2 localized to the mating projection tip (**Fig. 3A**, **Supplementary movie 3-4)** and their level started gradually decreasing ∼2-4 minutes before fusion of cells could be observed (**Fig. 3C**). These results demonstrate that Tpm1 and Tpm2 localize to the mating fusion focus and suggest a role for them in stabilizing formin-based actin cables during yeast mating. Previous studies have shown that the pheromone induced shmoo formation is dependent only on the formin Bni1 which is recruited via Bil2 and requires Bud6 activity^71,76^. So, we performed another experiment where we mated Δ*bnr1* cells expressing either mNG-^AS^Tpm1 or mNG-^AS^Tpm2 with wildtype cells of opposite mating type. We observed that both mNG-^AS^Tpm1 and mNG-^AS^Tpm2 localized to the mating projection tip with similar kinetics in Δ*bnr1* cells similar to wildtype cells **(Fig. 3B, 3D)**, suggesting that both Tpm1 and Tpm2 bind to Bni1-made actin filaments at the mating projection during the pheromone response in *S. cerevisiae.* These observations suggest a role for Tpm1 and Tpm2 in stabilization of Bni1-made actin cables during yeast mating and further strengthen our conclusion that Tpm1 and Tpm2 indiscriminately bind to formin-made filaments irrespective of the identity of the formin isoform in *S. cerevisiae*.

**Figure 3:**
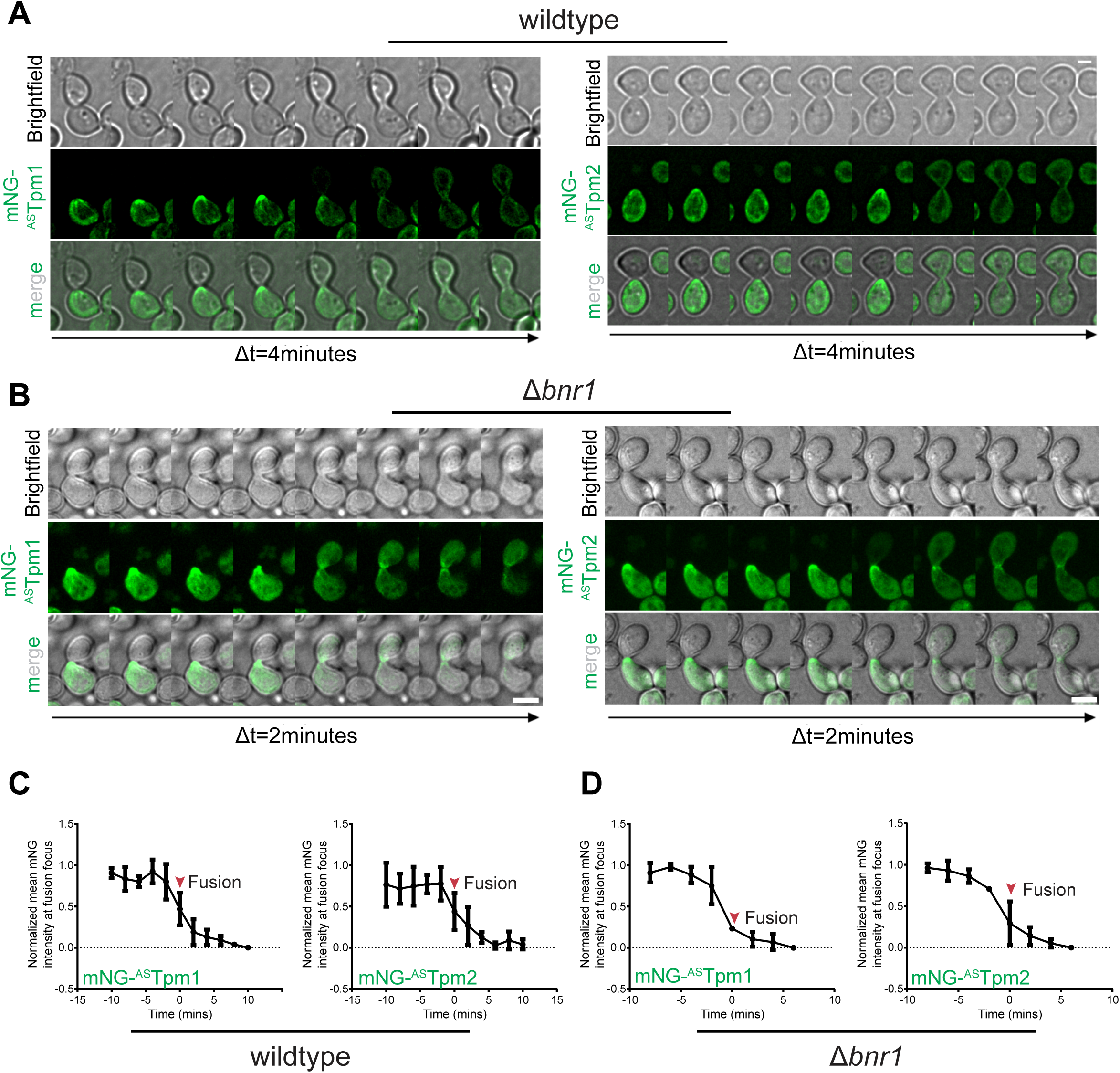
Tpm1 and Tpm2 co-localize at the fusion focus in mating yeast cells in a Bni1-de-pendent manner. **(A)** Representative time-lapse montages of haploid *MATa* type wildtype yeast cell expressing mNG-^AS^Tpm1 (left panel) or mNG-^AS^Tpm2 (right panel) mating with haploid *MATα* wildtype yeast cell; scale bar – 2μm. **(B)** Representative time-lapse montages of haploid *MATa* type ∆*bnr1* yeast cell expressing mNG-^AS^Tpm1 (left panel) or mNG-^AS^Tpm2 (right panel) mating with haploid *MATα* wildtype yeast cell; scale bar – 2μm. **(C-D)** Line plot showing protein accumulation kinetics of mNG-^AS^Tpm1 or mNG-^AS^Tpm2 at the mating projection tip in wildtype (C) or ∆*bnr1* (D) cells. t=0 rep-resents cell fusion (n=3 fusion events/strain).

### Tpm1 and Tpm2 protect equally well from cofilin-mediated actin filament severing

Tpm1 and Tpm2 isoforms are believed to play both shared and distinct roles in *S. cerevisiae*^16,39^. The major isoform Tpm is expressed 5-6 fold higher than Tpm2 and its absence leads to a near complete loss of actin cables^15,36,75^ **(Fig. S4B)**. A previous study reported that overexpression of Tpm2 could not restore actin cables in a Δ*tpm1* strain suggesting that Tpm2 and Tpm1 may have distinct functions^16^. We were intrigued by this observation as Tpm1 and Tpm2 have identical subcellular localization and dynamics as revealed by live-cell imaging of mNG-Tpm fusion proteins (**Fig. 1A**, **2G)**. Thus, we decided to revisit Tpm2 functionality and determine its subcellular roles in maintaining the actin cytoskeleton. Firstly, to test whether Tpm1 and Tpm2 impart similar protection against cofilin-mediated severing, we performed an in-vitro TIRF imaging^77,78^ with purified yeast tropomyosins (Tpm1, Tpm2), human Tpm1.7^79^ (positive control) and budding yeast Cofilin (Cof1) (**Fig. 4A**). We found that Tpm1.7, Tpm1 and Tpm2 coated-actin filaments displayed significantly lower severing events per unit time and length (**Fig. 4B**) and resulted in higher filament lengths after 2 min of cofilin addition as compared to uncoated control filaments (**Fig. 4C**). These observations suggest that Tpm1 and Tpm2 display similar biochemical activities of actin filament protection from severing action of cofilin, and hint towards an potential protective role of Tpm2 in cells.

**Figure 4:**
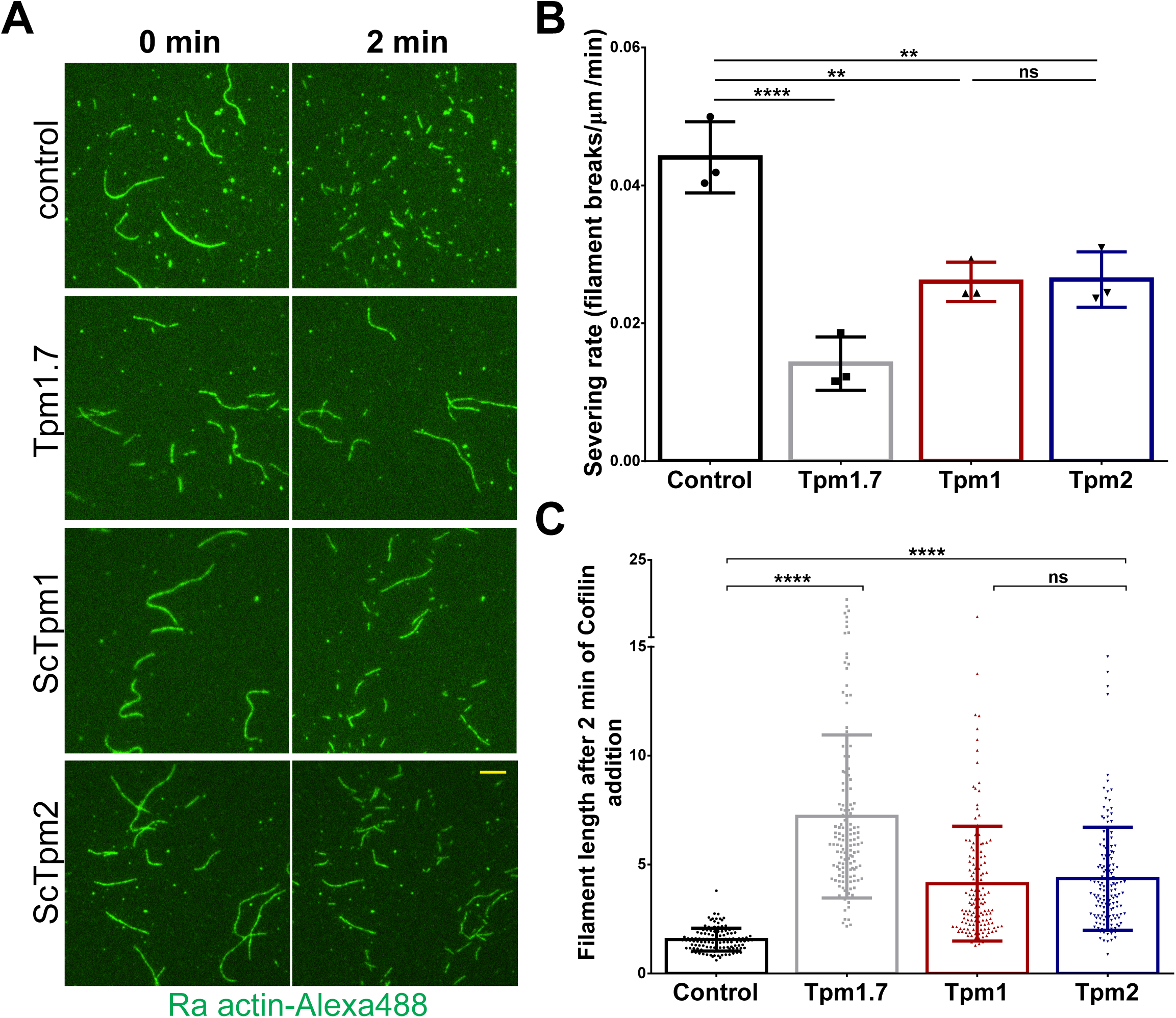
Tpm2 can protect actin filaments from cofilin-mediated severing *in vitro*. **(A)** Repre-sentative time-lapse images of the actin filaments before (left) and 2 min after the addition of recom-binant ScCofilin (right) in absence or presence of different Tpm isoforms; scale bar -10μm. **(B)** Plot representing number of severing events per μm of actin filaments per minute. Data shown for inde-pendent trials; n≥150 filaments in each condition, N=3. **(C)** Plot representing actin filament lengths after 2 minutes of addition of Cof1 to the samples in absence (control) or presence of specified tropo-myosin; n= ∼200 filaments, N=3. (Box represents 25^th^ and 75^th^ percentile, line represents median, whiskers represent minimum and maximum value; One-Way Anova with Tukey’s Multiple Comparisons test was used in (B) and (C) * p < 0.05, ** p < 0.01, *** p < 0.001, **** p < 0.0001)

### Tpm2 overexpression can compensate for loss of Tpm1 function *in vivo*

To coroborate our in-vitro findings and test whether Tpm2 could complement growth defects caused by loss of Tpm1, we overexpressed Tpm2 in a Δ*tpm1* strain using a high-copy number plasmid and surprisingly, found that Tpm2 overexpression could restore growth rates similar to wildtype levels in Δ*tpm1* cells **(Fig. S4A)**. This intriguing finding led us to investigate whether Tpm2 overexpression could also restore actin cables in Δ*tpm1* cells. Interestingly, we observed that Tpm2 overexpression could rescue actin cables similar in length and number to wildtype in ∼80% of Δ*tpm1* cells **(Fig. S4B, S4C)**. We confirmed Tpm2 overexpression using RT-qPCR data which revealed a ∼19 fold higher Tpm2 mRNA abundance after expression through high-copy number plasmid (*pRS425*-^pTpm2^*TPM2*) as compared to Tpm2 mRNA levels in wildtype and Δ*tpm1* cells **(Fig. S4D)**. This data clearly suggests that in contrary to previous understanding, elevated levels of Tpm2 can organize polarized actin cables in the absence of Tpm1, potentially by protecting actin cables from the severing action of cofilin. This represents a previously unrecognized activity for Tpm2 and raises further interesting questions about the shared and distinct roles of Tpm isoforms in *S. cerevisiae*.

### Mild increase in Tpm2 expression can restore actin cable organization in Δ*tpm1* cells

We assessed if increasing Tpm2 protein levels via low-copy plasmid expression under native Tpm1 and Tpm2 promoters (^pTpm1^*TPM1* and ^pTpm*2*^*TPM2*) could rescue defects of Δ*tpm1* cells. We tested their ability to rescue growth of Δ*tpm1* cells and found that both ^pTpm2^*TPM2* and ^pTpm1^*TPM2* constructs restored wildtype growth levels in Δ*tpm1* cells **(Fig. S4F)**. Expression of ^ptpm2^Tpm1 and ^pTpm1^*TPM1* (positive control) also restored normal growth levels in Δ*tpm1* cells **(Fig. S4F)**. F-actin staining with Alexa488-phalloidin showed rescue of actin cables by all Tpm2 and Tpm1 expression constructs in ∼90% Δ*tpm1* cells **(Fig. 5A, 5B)** to similar length as the wildtype (**Fig. 5C**). To determine the amount of Tpm2 mRNA after expression through low-copy plasmids, we performed RT-qPCR and found that Tpm2 mRNA levels showed ∼2 fold average increase when expressed under the pTpm1 promoter and did not show any detectable change from wildtype levels when expressed under its native *ptpm2* promoter **(Fig. S4E)**. Wildtype and Δ*tpm1* cells had similar Tpm2 mRNA levels suggesting Tpm2 expression is not sensitive to loss of Tpm1 **(Fig. S4F)**. In the absence of specific antibodies against Tpm1 and Tpm2, we used the functionality of mNG-^AS^Tpm fusion proteins to rescue phenotype in Δ*tpm1* cells and ascertain that increased protein levels of Tpm2 are reponsible for the rescue. High-copy plasmid (*pRS425*) expression of both mNG-^AS^Tpm1 and mNG-^AS^Tpm2 under their native promoters rescued the severe growth defects of Δ*tpm1* cells at 37°C but only mNG-^AS^Tpm1 and not mNG-^AS^Tpm2 rescued the defect when expressed from an integrated plasmid copy (*pRS305*) under their native promoters **(Fig.S5A)**. Quantification of mean mNG signal intensity per cell revealed that the highest expression was observed *in pRS425-*^pTpm1^*mNG-^AS^Tpm1* followed by *pRS425-*^pTpm2^*mNG-^AS^Tpm2*, *pRS305-*^pTpm1^*mNG-^AS^Tpm1* and pRS305-^pTpm2^mNG-^AS^Tpm2 **(Fig.S5B, S5C)**, consistent with our earlier observation that mNG-Tpm2 fusion is functional at higher expression levels as compared to native untagged Tpm2 **(Fig.S1E, S1F)**. We also performed western blotting with anti-mNG antibody to detect protein levels **(Fig. S5D)** and found that *pRS425-*^pTpm1^*mNG-^AS^Tpm1* had the highest expression levels and *pRS425-*^pTpm2^*mNG-^AS^Tpm2* and *pRS305-*^pTpm*1*^*mNG-^AS^Tpm1* only expressed to ∼20% and ∼10% of that level **(Fig. S5E).** Protein levels were not detectable in *pRS305-*^pTpm*2*^*mNG-^AS^Tpm2* containing Δ*tpm1* cells **(Fig. S5D)** and in conjunction with our mean intensity analysis **(Fig. S5B, S5C)** explain the lack of rescue observed in Δ*tpm1* cells containing *pRS305-*^pTpm*2*^*mNG-^AS^Tpm2*. Taken to-gether, these data clearly suggest that it is the increase in mNG-^AS^Tpm2 protein levels that lead to rescue of observed phenotype in Δ*tpm1* cells.

**Figure 5:**
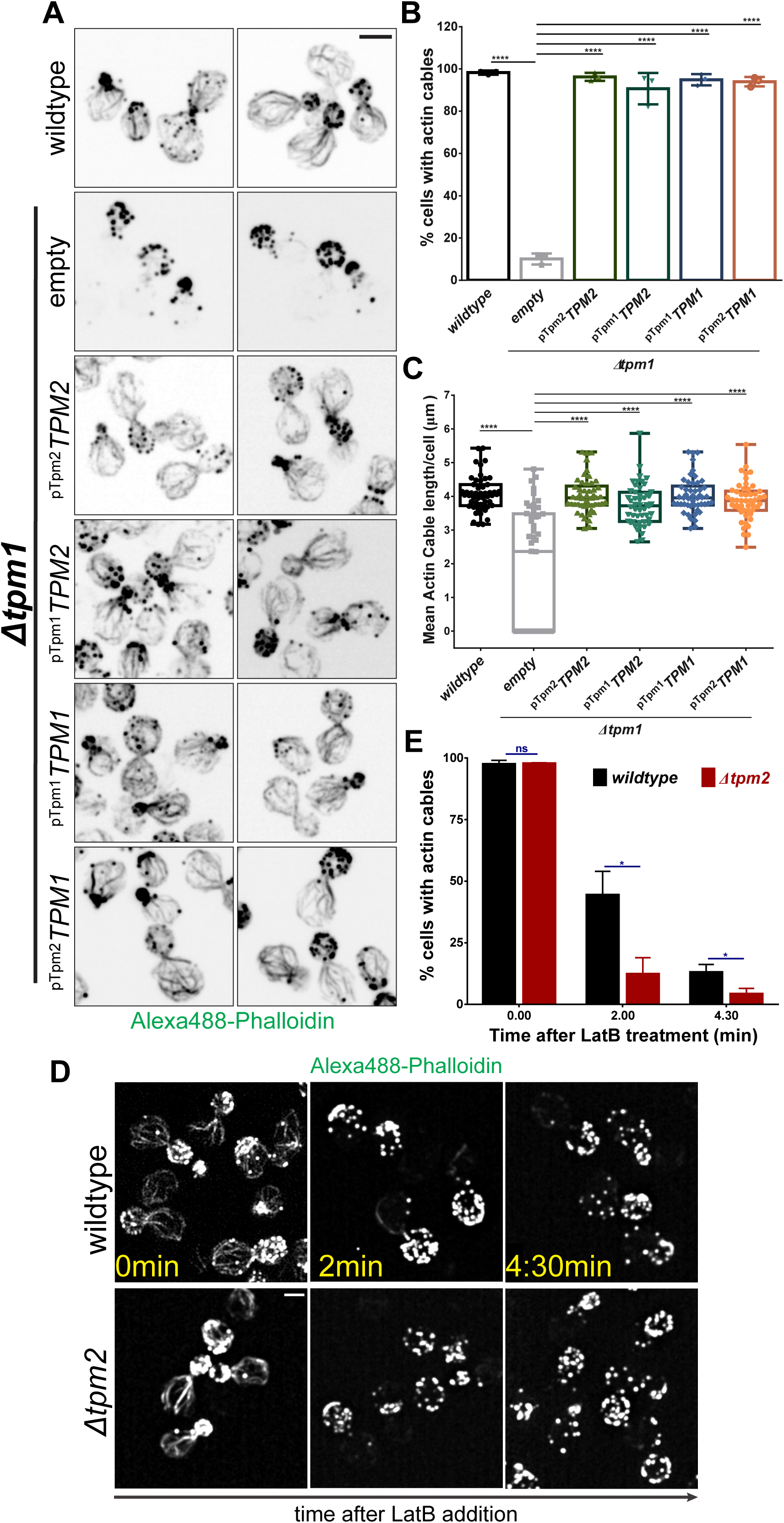
Increased Tpm2 expression can restore actin cables in Δ*tpm1* cells. **(A)** Representa-tive images of indicated yeast strains stained with Alexa488-phalloidin; scale bar – 3μm. **(B)** Plot representing mean percentage of cells with detectable actin cables in indicated yeast strains aver-aged over 3 biological replicates; n>150 cells for each strain per replicate, N=3. **(C)** Plot representing mean actin cable length per cell in the indicated yeast strains as per experiment in (A); n=50 cells for each strain. **(D)** Representative time-course images of wildtype and ∆*tpm2* cells treated with Latrun-culin B (67μM) and stained with Alexa488-phalloidin; scale bar -2μm. **(E)** Plot representing mean percentage of cells with detectable actin cables in indicated yeast strains averaged over 3 biological replicates; n>125 cells per strain per replicate, N=3. (Box represents 25^th^ and 75^th^ percentile, line represents median, whiskers represent minimum and maximum value; One-Way Anova with Tukey’s Multiple Comparisons test was used in (C) and Un-paired two-tailed t-test was used in (E), * p < 0.05, ** p < 0.01, *** p < 0.001, **** p < 0.0001)

Next, to test whether actin cable sensitivity is affected in cells lacking Tpm2, we assessed the presence of actin cables after treatment with the actin monomer sequestering drug, Latrunculin B (LatB) (**Fig. 5D**). We found that Δ*tpm2* cells are significantly more sensitive to the depolymerizing effect of LatB than wildtype cells **(Fig. 5D, 5E)**, suggesting that Tpm2 does play previously unrecognized detectable roles in actin cable stability in conditions of stress and contributes to robustness of the actin cable network.

Together, these results show that even minor increases in Tpm2 expression can restore normal actin cable organization in Δ*tpm1* cells suggesting that Tpm2 protein is present at limiting levels but retains the function of actin cable protection from severing factors like cofilin *in vivo* and *in vitro*. Tpm2 can, therefore, act as a safeguard mechanism for cells to protect against defects in Tpm1, providing functional redundancy and robustness against cellular stressors for actin cable maintenance in *S. cerevisiae*.

### Tpm2 can solely organize a functional actin cable network in *S. cerevisiae*

Actin cables in *S. cerevisiae* are required for functions such as anterograde transport of vesicles and organelles^22,36,41,64,65,75^, maintenance of mitochondrial inheritance and morphology^53,54,80,81^, and clearance of protein aggregates from the bud^82,83^ while retaining damaged mitochondria in the mother via retrograde actin cable flow (RACF)^40,84^. We asked whether Tpm2 organized actin cables could perform all these functions and if Tpm1-organized and Tpm2-organized actin cable networks had functional differences. We systematically analyzed these functional readouts using fluorescent reporters in wildtype, Δ*tpm1*, and Δ*tpm1* cells expressing Tpm2 in low-copy number and high-copy number plasmids. We used exogenous expressed ymScarleti-Sec4^36^ integrated into the *ura3/leu2* locus to assess vesicle targeting to the bud via Myo2-dependent transport on actin cables^22^ (**Fig. 6A**, **S6A)**. The ratio of bud-to-mother Sec4 fluorescence was calculated for all strains and the results showed that vesicle targeting to the bud was restored to levels similar to wildtype cells by expression of both ^pTpm2^*TPM2* and ^pTpm1^*TPM2* constructs (**Fig. 6C**, **S6C)**. Tpm1 expression also rescued vesicle targeting as a positive control under pTpm1 and pTpm2 promoters **(6A, 6C, S6C)**. These results suggest that both Tpm2 and Tpm1 can individually facilitate normal anterograde flow of cargo and vesicles to the growing bud via facilitation of Myo2 processivity on actin cables, in agreement with previous *in vitro* experiments where both Tpm1 and Tpm2 increased processivity of the type-V myosin, Myo2, on coated actin filaments^41^.

**Figure 6:**
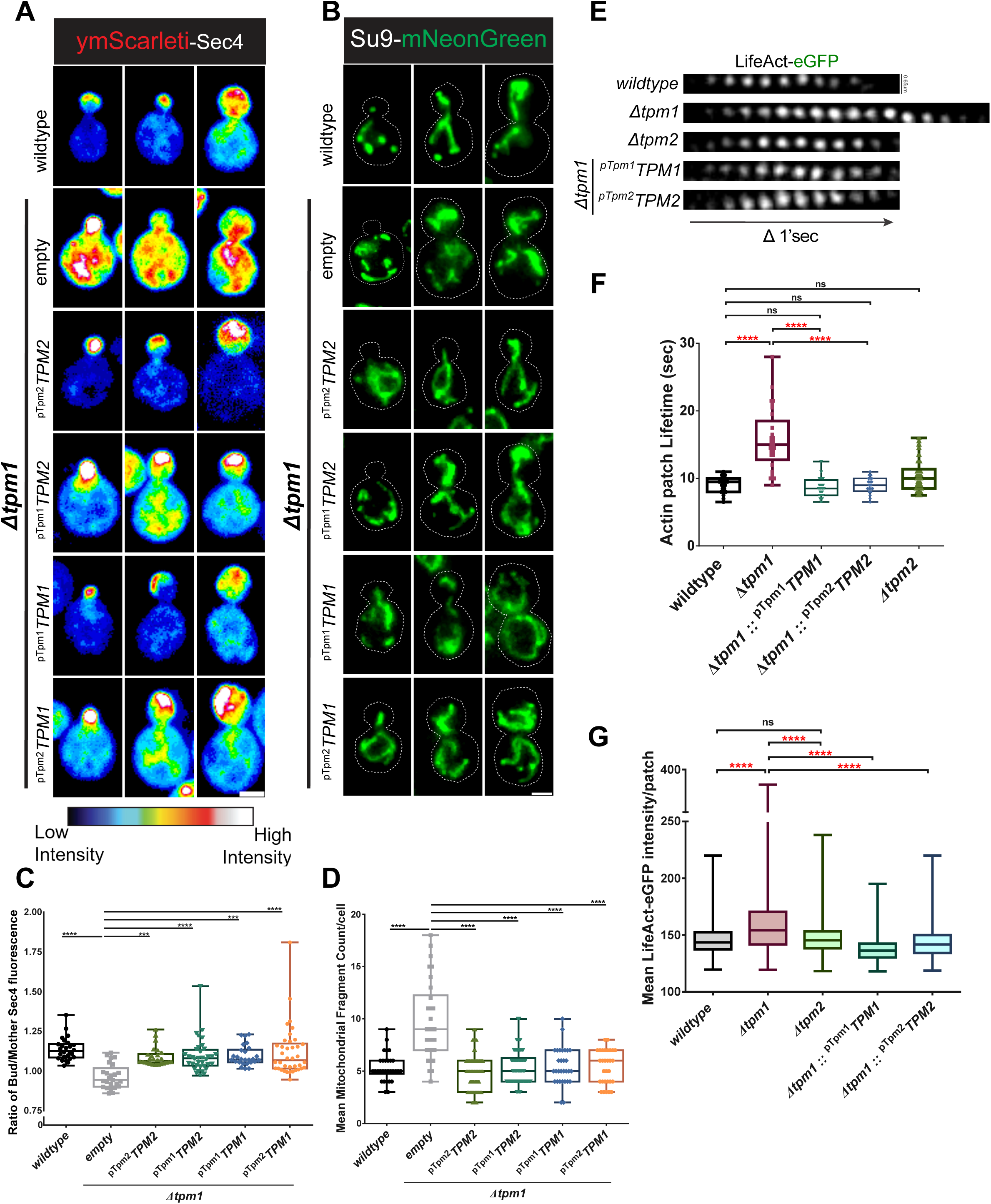
Tpm2 can independently organize a functional actin cytoskeleton in Δ*tpm1* cells and. **(A)** Representative images of indicated yeast strains expressing ymScarleti-Sec4; scale bar – 3μm. **(B)** Representative images of indicated yeast strains expressing Su9-mNeonGreen; Scale bar – 3μm. **(C)** Plot representing ratio of bud/mother ymScarleti-Sec4 fluorescence per cell in the indicated yeast strains; n≥25 cells per strain. **(D)** Plot representing mean mitochondrial fragment count per cell in the indicated yeast strains; n≥30 cells per strain. **(E)** Representative time-lapse images of actin patch dynamics in cells with indicated genotypes expressing LifeAct-eGFP, scale bar – 0.65μm. **(F)** Box and Whiskers plot showing actin patch lifetimes in the indicates yeast strains, (n>20 patches/strain). **(G)** Box and Whiskers plot showing mean LifeAct-eGFP fluorescence in the indicates yeast strains, (n≥735 cells/strain). (Box represents 25th and 75th percentile, line represents median, whiskers represent minimum and maximum value; One-Way Anova with Tukey’s Multiple Comparisons test was used in (C), Kruskal-Wallis test with Dunn’s multiple comparisons test was used in (D), (G), and (F); * p < 0.05, ** p < 0.01 * p < 0.05, ** p < 0.01, *** p < 0.001, **** p < 0.0001)

Next, we assessed whether the hyper-fragmented mitochondrial morphology, a characteristic of Δ*tpm1* cells^53,54^, could be restored by exogenous expression of Tpm2. For visualization of mitochondrial morphology, we used Su9-mNeonGreen^85^ (subunit 9 of the F1-F0 ATPase fused to mNeon-Green) construct integrated at the *leu2/ura3* locus and quantified average mitochondrial fragment number per cell (**Fig. 6B**, **S6B)**. Δ*tpm1* cells showed increased mean mitochondrial fragment number as compared to wildtype while expression of Tpm2 or Tpm1 under native promoters in low copy or high copy plasmids restored the average mitochondrial fragment number to wildtype levels (**Fig. 6D**, **S6D)**. These results suggest that Tpm2-bound actin cables can interact and crosstalk normally with mitochondria to maintain its normal morphology. The underlying mechanisms of how actin cables control mitochondrial morphology in budding yeast remain largely unknown and future studies are required to understand this enigmatic phenomenon.

Tpm2 has been previously implicated in negative regulation of retrograde actin cable flow (RACF) rates, possibly via its inhibitory effect on binding of the type-II myosin (Myo1) to actin filaments^39,40^. We independently replicated this experiment using LifeAct-eGFP^69,86^ as an actin cable marker and corroborated that RACF rates are increased in the absence of Tpm2 and exogenous expression of Tpm2 and not Tpm1 in Δ*tpm2* cells restores RACF rates to wildtype levels **(Fig. S6E, S6F)**. RACF rates were similar to wildtype in Δ*tpm1* cells in which Tpm2 was expressed to restore actin cable suggesting again that loss of Tpm1 did not affect RACF rates in cells **(Fig. S6F)**. These observations suggest that Tpm2 has gained distinct regulatory function from Tpm1 in negatively regulating RACF while retaining its actin protective activity.

### Global Tropomyosin levels are important for proper function and homeostasis between linear and branched actin in *S. cerevisiae*

Homeostasis between linear and actin networks in cells is maintained by various actin-binding proteins that coordinate and compete to generate actin filament network diversity in cells^87–91^. In yeast, formin proteins nucleate and maintain linear actin filaments which bundle and form cables^64,65^ while Arp2/3 complex generates branched actin filaments which form a dense actin network at sites of endocytosis called actin patches^92,93^. Actin cables and patches interplay with each other while competing for the available cytoplasmic pool of actin monomers^88,90,94,95^. Tpm1 and Tpm2 localize to actin cables (**Fig. 1A**) and are majorly excluded from actin patches due to the activity of fimbrin (Sac6) which majorly localizes to actin patches^52,87^. We have previously shown that Tpm1 and Tpm2 localize to actin patches in ∆*sac6* cells^44^ and a recent study observed similar targeting of Tpm to actin patches in the absence of Capping protein (Cap1/2 heterodimer) suggesting a tight regulation of protein composition between these two networks^89^. Tpm loss or overexpression has been shown to opposingly affect total number of actin patches^90^. We asked whether loss of actin cables observed in Δ*tpm1* cells could lead to effects on actin patch-mediated endocytosis. We used LifeAct-eGFP as a marker for actin patches (**Fig. 6E**) and observed that actin patch lifetime was significantly increased in Δ*tpm1* cells (∼15 sec) as compared to wildtype and Δ*tpm2* cells (∼10 sec) **(Fig. 6E, 6F)**. The increased patch lifetime was restored to wildtype levels upon exogenous expression of either Tpm1 or Tpm2 from a low-copy plasmid **(Fig. 6E, 6F)**, suggesting that the observed increase is caused due to the global decrease in Tpm levels and loss of actin cables, and not specifically due to loss of Tpm1. We next tested whether the loss of actin cables observed in Δ*tpm1* cells lead to increased accumulation of actin at the actin patches and found that mean LifeAct-eGFP intensity at actin patches was higher in Δ*tpm1* cells as compared to wildtype and Δ*tpm2* cells (**Fig. 6G**). The mean intensity was restored to wildtype levels upon exogenous expression of Tpm1 and Tpm2 in Δ*tpm1* cells (**Fig. 6G**), suggesting that reduction in global Tpm levels leads to dysregulation of actin cables, increased actin accumulation at the actin patches, eventually leading to abnormal endocytosis dynamics. These results highlight the isoform independent importance of Tpm in maintaining normal balance and homeostasis between actin networks in a common cytoplasmic volume in *S. cerevisiae* and opens avenues for future study of actin network crosstalk via effects of actin-binding protein in yeast and other eukaryotes.

Overall, our results suggest that Tpm2 can independently organize a functional actin cytoskeleton in *S. cerevisiae*. Our work highlights a previously unappreciated function of Tpm2 and demonstrates functional redundancy between Tpm isoforms despite evolutionary divergence and evolution of distinct functions such as RACF regulation. This functional redundancy may benefit cells under stress conditions and contribute to robust cell survival and viability.

## Discussion

### Making Functional Tropomyosin fusion proteins

In this study, we characterize and present functional fluorescently tagged-tropomyosin fusion proteins which make it possible to probe tropomyosin isoform localization, functions, and competition with other actin-binding proteins in live cells. Building on our previous work^44^, we have shown that mNG-Tpm1 and mNG-Tpm2 are functional and can restore viability of Δ*tpm1*Δ*tpm2* cells which display synthetic lethality **(Fig. S1E)**. To our knowledge, this represents the first demonstration and characterization of tagged tropomyosin proteins which clearly report localization while near completely restoring growth and actin organization in cells lacking native Tpms (**Fig. 1A-D**). Previous work in *S. pombe* had also identified N-terminus of *S. pombe* Tpm (Cdc8) as being favorable for tagging while the C-terminal fusion rendered the protein completely non-functional^46^. A key difference in our study is the presence of a 40-amino acid linker between the fluorescent tag and the Tpm protein^44^, which seems to enhance functionality and allow proper head-to-tail contacts between Tpm dimers, as evident by the clear localization observed on actin cables which is not seen in case of direct N-terminal tagging of Tpm^96^. N-terminal tagging also has its caveats as it blocks N-terminal acetylation of Tpm which is essential for normal Tpm binding to F-actin in cells^47,48^ and also for regulation of other actin-binding proteins such as myosin-II and myosin-V^24,45^. To overcome this limitation, we generated mNG-^AS^Tpm fusion proteins which contain an Alanine-Serine (-AS-) dipeptide to mimic the effect of N-terminal acetylation. The use of -AS-dipeptide is routinely used to purify Tpm proteins from *E. coli* as it restores normal binding affinity^50^ and its addition to the N-terminus of Tpm in the mNG-Tpm fusion improved their functionality by restoring normal length of actin cables in our experiments (**Fig. 1B**, **1C)**. The observed effect may be due to increased binding affinity of -AS-containing Tpm for F-actin in cells as compared to unacetylated tagged Tpm and also suggests a role for Tpm activity in cell size dependent scaling of actin cables in yeast. Previous work suggested that acetylation decreases the destabilizing effect of the positive charge of the N-terminus on the coiled-coil ‘a’ position where a hydrophobic residue normally resides^50^. A later study proposed that the -AS-dipeptide ensures that the Met1 stays in an alpha-helical structure by introducing stabilizing hydrogen bonds between the Ser and Met residues^97^. Thus, -AS-dipeptide addition likely does not mimic N-terminal acetylation structurally but acts via maintaining N-terminus stability. Future work is required to understand in detail the effects of the linker and -AS-dipeptide on Tpm biochemical activity and effect on interactions with other actin-binding proteins (ABPs).

### Spatial sorting of Tpm isoforms

The molecular basis of Tpm isoform sorting has remained an unresolved enigma in the field. Various studies over the years have implicated different mechanisms for spatial sorting of Tpm isoforms across different model systems. While *S. pombe* Tpm (Cdc8) exhibits sorting based on preference between N-terminal acetylation status and formin isoform^45^, mammalian Tpms have been proposed to sort through either formin-based nucleation^32^ or relative concentration dependent mechanisms^30^. However, N-terminal acetylation did not bias localization of Cdc8 to filaments made by a particular *S. pombe* formin isoform in a minimal in-vitro reconstituted set-up^98^, in contrast to the earlier in vivo observations^45^, suggesting that interaction of the distinct Cdc8 forms (unacetylated vs acetylated) with distinct formins may not be enough for the spatial sorting observed and may require additional cellular factors. In this study, we utilized our functional mNG-Tpm constructs to address this long-standing question of Tpm isoform sorting in the model *S. cerevisiae* and find a formin-independent mode of binding of Tpm isoforms to actin cables, in contrast to the formin-dependent modes observed in *S. pombe*^31^. We also performed first-ever dual-color imaging of Tpm1 and Tpm2 simultaneously in live cells which highlights the multiplexing possibilities of our tagging strategy to study Tpm isoform diversity (**Fig. 2G**). Strikingly, while *tpm1* shows distinct genetic interactions with formins Bnr1 and Bni1^36^, it did not show any preference for filaments made by any one of these isoforms in our study. This seemingly contrasting observations could be explained by the observations from *Shin et. al., 2018*^99^, where difference in activities of Bnr1 and Bni1 in the presence of limiting levels of Tpm2 inside cells could explain why loss of Bnr1 or Bni1 in Δ*tpm1* cells has opposite effects on cell growth and fitness. Development of probes enabling measurement of formin activity in cells could help confirm this hypothesis in the future. While it is possible that N-terminal tagging may influence formin-based Tpm isoform sorting, C-terminally tagged constructs cannot currently be used to address this question due to their non-functional nature. Thus, there is still scope for developments and improved strategies for Tpm visualization to revisit these interesting questions. Overall, our data suggests distinct evolutionary adaptations and strategies for Tpm isoform sorting across species, which may be contingent on species-specific needs and adaptations.

### Shared and Distinct functions of Tpm1 and Tpm2

Our work also addresses the question of functional overlap and divergence of Tpm isoforms where we report a previously unknown role of the “minor” isoform Tpm2 in actin cable protection suggesting that Tpm2 has retained its actin protective function despite evolutionary divergence and gain of distinct regulatory activity in controlling retrograde actin cable flow as compared to Tpm1^39^. We find that Tpm1 and Tpm2 protect equally well from cofilin *in vitro* (**Fig. 4D-F**), suggesting that their binding sites on F-actin may overlap with cofilin-binding site. However, their distinct regulation of Retrograde Actin cable flow (RACF) which is controlled by myosin-II (Myo1) suggests distinct effects on myosin-II binding possibly through different overlap with myosin-II binding site on the actin filament^39^. Structural data about their binding will reveal more insights into these possibilities. A possible reason why the earlier study could not complement Δ*tpm1* cells with Tpm2 is the use of galactose-induction based overexpression from a high-copy plasmid which would result in abnormally high levels of Tpm2 in cells, causing toxicity and cell death^16^. From our results, it’s clear that Tpm2 is present at limiting levels and a mild increase in expression restores normal growth and actin cytoskeleton in Δ*tpm1* cells (**Fig. 4A-C**). The reason to keep such low levels might be to maintain Tpm2 specific function in RACF-mediated asymmetric inheritance of damaged organelles and protein aggregates between mother and bud^39,40,82,84^, while also acting as a latent reservoir for actin cable maintenance in stressed conditions. Consistent with this, we found cells lacking Tpm2 show decreased cable stability in presence of the actin depolymerizing drug LatB **(Fig. 4G, 4H)**. Our data suggests that such functional redundancy among diverged Tpm isoforms may confer robustness to actin-dependent processes and increase chances of cell survival under certain stressed conditions. In the future, it remains to be understood whether structural differences in F-actin binding modes^34^ or other factors such as post-translational modifications^25,48^ in conjunction with expression levels and spatio-temporal regulation, etc. contribute to the shared and distinct roles of closely related Tpm isoforms in *S. cerevisiae* and other eukaryotes.

### Tropomyosin levels control global actin regulation and cross-talk of actin networks

Actin monomeric pool in the cell is used to maintain diverse actin filament network types that differ in geometry, composition of actin-binding proteins (ABPs), turnover and thus, their function^88^. Tropomyosin majorly binds to linear actin filaments nucleated by formin proteins and prevents branching of linear actin filaments by inhibiting Arp2/3 complex-mediated branch nucleation on existing filaments^88,98,99^. Linear actin cables and branched actin patches are the two major actin networks in yeast with Tpm majorly localizing to the cables. Our data suggest that global decrease in Tpm levels not only cause a loss of actin cables but also affect actin accumulation and dynamics at actin patches which are sites of endocytosis (**Fig. 5E-G**). This demonstrates that Tpm1 and Tpm2 levels are important for global actin network homeostasis and suggests that dysregulation of actin cables caused by decreased Tpm levels trigger dysregulation of other actin networks which may contribute to cellular defects observed in ∆*tpm1* cells. It remains to be addressed whether these effects are entirely due to increased actin accumulation or also accompany changes in ABP composition of the branched actin network. Future work with emphasis on network composition is required to delineate exact molecular basis of such effects in cells with loss of normal actin network homeostasis. Our study, thus, opens exciting avenues to understand effects of ABPs on actin networks at a systems level in the future.

## Materials and Methods

### Plasmid construction

All plasmids used in this study are listed in Supplementary Table 1. Plasmids were constructed by assembling DNA fragments amplified by PCR into restriction endonuclease-digested vectors using NEB Hifi Builder (NewEngland BioLabs; cat.no: E2621L). The plasmids were confirmed with re-striction digestion and sequencing.

### Yeast Strains construction

All yeast strains used in this study are listed in Supplementary Table 2. S288c genetic background was used for all modifications. Yeast transformation was done using the Lithium Acetate based method as previously described^102^. Deletion and tagging of genes was done using the homologous recombination of PCR cassettes as described previously^103^. The strains were confirmed using colony PCR and fluorescence microscopy.

### Live-cell Imaging of tropomyosin isoforms

Yeast strains were grown overnight in Synthetic Complete (SC) Media at 25°C. The overnight culture was used to inoculate a secondary culture which was allowed to grow till mid-log phase. The cells were then adhered on a Concanavalin A-coated glass-bottom dish (Cellvis cat: no: D35C4-20-1.5-N) containing SC media. Images were acquired using the Andor Dragonfly 502 spinning disk confocal system (Oxford Instruments) equipped with a fully-motorized Leica Dmi8 inverted microscope setup. Z-stacks were acquired using a 100x oil objective, captured with the Andor Sona scMOS camera and deconvolved using Andor Fusion software. The images were processed using Fiji and quantified using a previously described protocol^56,104^.

### Actin cable staining with phalloidin

In order to visualize the actin cables, cells were stained with Alexa488/Rhodamine Phalloidin and imaged using a previously described protocol^56,104^. Briefly, cells were grown at 23℃ until early-mid log phase in Yeast extract/Peptone/2%Glucose (YPD) media. The cells were then fixed twice with 4% paraformaldehyde in YPD and 4% paraformaldehyde in 1x PBS for 45 minutes each and washed thrice with 1x PBS. Rhodamine/Alexa488 phalloidin (Invitrogen cat: no: R415 and cat: no: A12379 respectively) was added to a final concentration of 0.4µM and incubated overnight with rotation at 4℃. Next day, the cells were washed thrice with 1X PBS. The cells were adhered on 6% Concanavalin A (Sigma cat: no: C2010) coated glass bottom dishes and z-stacks (step size = o.2μm, 31 slices) were acquired using Andor Dragonfly 502 Spinning Disk system and deconvolved using Andor Fusion Software.

### Actin and Tropomyosin Cable quantification

Actin/Tropomyosin cable length and number was done exactly as previously described^104^. An example of the quantification procedure and workflow are shown in **Figure. S7A**.

### Yeast mating experiment

Yeast strains of opposite mating types were grown overnight at 25°C and mixed in equal ratio. The mixture of the two strains were seeded on a 1.2% Agarose pad or on a 6% Concanavalin-A coated glass bottom dish and allowed to grow at 25°C for 30-45 minutes. Time-lapse images were acquired at an interval of 2 min to capture mating events.

### Purification and labeling of actin

Rabbit skeletal muscle actin was purified from acetone powder generated from frozen ground hind leg muscle tissue of young rabbits (PelFreez, USA). Lyophilized acetone powder stored at −80 °C was mechanically sheared in a coffee grinder, resuspended in G-buffer (5 mM Tris-HCl pH 7.5, 0.5 mM Dithiothreitol (DTT), 0.2 mM ATP, and 0.1 mM CaCl_2_), and cleared by centrifugation for 20 min at 50,000 × g. The supernatant was collected and further filtered with the Whatman paper. Actin was then polymerized overnight at 4 °C, slowly stirring, by the addition of 2 mM MgCl_2_ and 50 mM NaCl to the filtrate. Next morning, NaCl powder was added to a final concentration of 0.6 M, and stirring was continued for another 30 min at 4 °C. F-actin was pelleted by centrifugation for 2.5 hrs at 280,000 × g. The pellet was solubilized by dounce homogenization and dialyzed against G-buffer for 48 h at 4 °C. Monomeric actin was precleared at 435,000 × g and loaded onto a Sephacryl S-200 16/60 gel-filtration column (Cytiva, USA) equilibrated in G-Buffer. Fractions containing actin were stored at 4 °C. To biotinylate actin, purified G-actin was first dialyzed overnight at 4 °C against G-buffer lacking DTT. The monomeric actin was then polymerized by the addition of an equal volume of 2X labeling buffer (50 mM imidazole pH 7.5, 200 mM KCl, 0.3 mM ATP, 4 mM MgCl_2_). F-actin was then mixed with a fivefold molar excess of NHS-XX-Biotin (Merck KGaA, Germany) and incubated in the dark for 15 h at 4 °C. F-actin was pelleted by centrifugation at 450,000 × g for 40 min at room temperature, The pellet was rinsed with G-buffer, homogenized with a dounce and dialyzed against G-buffer for 48 h at 4 °C. Biotinylated monomeric actin was purified further on a Sephacryl S-200 16/60 gel-filtration column as above. Aliquots of biotin-actin were snap-frozen in liquid N_2_ and stored at −80 °C.

To fluorescently label actin, G-actin was polymerized by dialyzing overnight against modified F-buffer (20 mM PIPES pH 6.9, 0.2 mM CaCl_2_, 0.2 mM ATP, and 100 mM KCl). F-actin was incubated for 2 h at room temperature with a fivefold molar excess of Alexa-488 NHS ester dye (Thermo Fisher Scientific, USA, cat: no: A20100). F-actin was pelleted by centrifugation at 450,000 × g for 40 min at room temperature, and the pellet was resuspended in G-buffer, homogenized with a dounce and further incubated on ice for 2 h to depolymerize the filaments. The monomeric actin was then re-polymerized on ice for 1 h by the addition of 100 mM KCl and 1 mM MgCl_2_. F-actin was once again pelleted by centrifugation for 40 min at 450,000 × g at 4 °C. The pellet was homogenized with a dounce and dialyzed overnight at 4 °C against 1 L of G-buffer. Next morning, the solution was precleared by centrifugation at 450,000 × g for 40 min at 4 °C. The supernatant was collected, and concentration and labeling efficiency of actin were determined. Labelled actin was stored at 4 °C.

### Purification of profilin

Human profilin-1 was expressed in *E. coli* strain BL21 (pRare) to log phase in LB broth at 37 °C and induced with 1 mM IPTG for 3 h at 37 °C. Cells were then harvested by centrifugation at 15,000 × g at 4 °C and stored at −80 °C. For purification, pellets were thawed and resuspended in 30 mL lysis buffer (50 mM Tris-HCl pH 8, 1 mM DTT, 1 mM PMSF and protease inhibitors (0.5 μM each of pepstatin A, antipain, leupeptin, aprotinin, and chymostatin), and the solution was sonicated on ice by a tip sonicator. The lysate was centrifuged for 45 min at 120,000 × g at 4 °C. The supernatant was then passed over 20 ml of Poly-L-proline conjugated beads in a disposable column (Bio-Rad, USA). The beads were first washed at room temperature in wash buffer (10 mM Tris pH 8, 150 mM NaCl, 1 mM EDTA, and 1 mM DTT) and then washed again with two column volumes of 10 mM Tris pH 8, 150 mM NaCl, 1 mM EDTA, 1 mM DTT, and 3 M urea. Protein was then eluted with five column volumes of 10 mM Tris pH 8, 150 mM NaCl, 1 mM EDTA, 1 mM DTT, and 8 M urea. Pooled and concentrated fractions were then dialyzed in 4 L of 2 mM Tris pH 8, 0.2 mM EGTA, 1 mM DTT, and 0.01% NaN3 for 4 h at 4 °C. The dialysis buffer was replaced with fresh 4 L buffer, and the dialysis was continued overnight at 4 °C. The protein was centrifuged for 45 min at 450,000 × g at 4 °C, concentrated, aliquoted, flash-frozen in liquid N2, and stored at −80 °C.

### Purification of yeast and human Tpm isoforms

All budding yeast and human tropomyosins were expressed in *E. coli* strain BL21 to log phase in LB broth at 37 °C and induced with 1 mM IPTG for 3 h at 37 °C. Cells were then harvested by centrifugation at 15,000 × g at 4 °C and stored at −80 °C. For purification, pellets were thawed and resuspended in 30 mL lysis buffer (20 mM Tris-HCl pH 7.5, 0.5 M NaCl, 5 mM MgCl_2_, 1 mM PMSF, protease inhibitors (0.5 μM each of pepstatin A, antipain, leupeptin, aprotinin, and chymostatin)), and the solution was sonicated on ice by a tip sonicator. The lysate was incubated at 80°C for 10 min in a water bath and at room temperature for another 10 min. The lysate was centrifuged for 30 min at 30,000 x g at 4 °C, and the pellet was discarded. The protein was precipitated by adding 0.3M HCl to the supernatant till the pH reached 4.7. The solution was centrifuged for 30 min at 30,000 x g at 4°C. The pellet was resuspended in 25 mL of wash buffer (100 mM Tris-HCl pH 7.5, 0.5 M NaCl, 5 mM MgCl_2_, 1 mM DTT). The acid precipitation and accompanying centrifugation step were repeated again. The pellet was then resuspended in 25 mL of wash buffer and dialyzed in 2 L of dialysis buffer (20 mM HEPES pH 6.8, 50 mM NaCl, 0.5 mM DTT) overnight at 4 °C. The dialyzed solution was loaded to a 5 ml HiTrap Q HP column (Cytiva). The protein fractions were eluted with a linear gradient of NaCl (50– 600 mM). The fractions were analyzed by SDS-PAGE to determine the ones containing Tpm. They were concentrated and further loaded on a Superose 6 gel-filtration column (Cytiva) pre-equilibrated with 20 mM Tris (pH 7.5), 50 mM KCl, 2 mM MgCl2, 1 mM DTT. Peak fractions were collected, con-centrated, aliquoted, and flash-frozen in liquid N2 and stored at −80 °C.

### Purification of yeast cofilin

His-tagged yeast Cof1 was expressed in *E. coli* strain BL21 to log phase in LB broth at 37 °C and induced with 1 mM IPTG for 3 h at 37 °C. Cells were then harvested by centrifugation at 15,000 × g at 4 °C and stored at −80 °C. For purification, pellets were thawed and resuspended in 30 mL lysis buffer (50 mM phosphate buffer pH 8, 20 mM Imidazole, 0.3 M NaCl, 1 mM PMSF, protease inhibi-tors), and the solution was sonicated on ice by a tip sonicator. The lysate was centrifuged for 45 min at 120,000 × g at 4 °C. The supernatant was incubated with 2 mL of washed Ni-NTA beads for 2 hrs at 4 °C. The solution was centrifuged at 1000 xg for 5 minutes. The supernatant was removed. The beads were washed 3 times with 10 mL of wash buffer (50 mM Phosphate buffer pH 8, 20 mM Imid-azole, 0.3 M NaCl, 1 mM DTT) and subsequently centrifuged at 1000 xg for 2 min each time to remove the supernatant. Protein was then eluted with 1 mL of elution buffer (50 mM phosphate buffer pH 8, 250 mM Imidazole, 0.3 M NaCl, 1 mM DTT). Pooled and concentrated fractions were then loaded on a Superose 6 gel-filtration column (Cytiva) pre-equilibrated with 20 mM HEPES pH 7.5, 50 mM KCl, 0.5 mM DTT. Peak fractions were collected, concentrated, aliquoted, and flash-frozen in liquid N2 and stored at −80 °C.

### TIRF microscopy

Glass coverslips (60 × 24 mm; Thermo Fisher Scientific, USA) were first cleaned by sonication in detergent for 20 min, followed by successive sonications in 1 M KOH, 1 M HCl, and ethanol for 20 min each. Coverslips were then washed extensively with H_2_O and dried in an N_2_ stream. The cleaned coverslips were coated with 2 mg/mL methoxy-polyethylene glycol (mPEG)-silane MW 2000 and 2 µg/mL biotin-PEG-silane MW 3400 (Laysan Bio, USA) in 80% ethanol (pH 2.0) and incubated over-night at 70 °C. Flow cells were assembled by rinsing PEG-coated coverslips with water, drying with N_2_, and adhering to μ-Slide VI0.1 (0.1 mm × 17 mm × 1 mm) flow chambers (Ibidi, Germany) with double-sided tape (2.5 cm × 2 mm × 120 μm) and epoxy resin for 5 min (Devcon, USA). Before each reaction, the flow cell was sequentially incubated for 1 min each with 4 μg/ml streptavidin and 1% BSA in 20 mM HEPES pH 7.5 and 50 mM KCl. The flow cell was then equilibrated with TIRF buffer (10 mM imidazole, pH 7.4, 50 mM KCl, 1 mM MgCl_2_, 1 mM EGTA, 0.2 mM ATP, 10 mM DTT, 2 mM DABCO, and 0.5% methylcellulose [4000 cP]).

Actin filaments were assembled by introducing 1 µM 15% Alexa 488 labelled 0.1% biotinylated actin and 2 µM profilin in the flow cell, with or without 10 μM Tpm. Filaments were allowed to grow for 2 to 3 min. For experiments containing Tpm, the flow cell was then rinsed with TIRF buffer supplemented with 10 μM Tpm to remove free actin. The solution was then replaced with 10 μM Tpm and 150 nM Cof1. For control experiments containing no Tpm, both washing and cofilin addition step had no Tpm in the solution.

### Image acquisition and analysis

Single-wavelength time-lapse TIRF imaging was performed on a Nikon-Ti2000 inverted microscope equipped with a 40 mW Argon laser, a 60X TIRF-objective with a numerical aperture of 1.49 (Nikon Instruments Inc., USA), and an IXON LIFE 888 EMCCD camera (Andor Ixon, UK). One pixel was equivalent to 144 × 144 nm. Focus was maintained by the Perfect Focus system (Nikon Instruments Inc., Japan). Time-lapse images were acquired every 2 s using Nikon Elements imaging software (Nikon Instruments Inc., Japan). The sample was excited by a 488 nm laser for imaging. Images were analyzed in Fiji. Normalized rates of severing by cofilin, in the presence or absence of different Tpm isoforms, were determined by counting the number of severing events as a function of time in a field of view divided by the total length of all filaments in the field of view. The average filament length due to severing was measured ∼2 min after the cofilin flow-in. Data analysis and curve fitting was carried out in Microcal Origin.

### RNA extraction and RT-qPCR

Hot phenol method was used to extract RNA from yeast cells^105^. Overnight grown yeast cells were diluted and allowed to grow until OD at 600nm reached 1. Cells were treated with equal volumes of TES and water saturated acidic phenol and precipitated at -80℃ to extract RNA. The extracted RNA was then treated with DNAse A and this was used as a template for first strand cDNA synthesis. Following this, qRT-PCR was set up using the cDNA to check the expression levels. Gene specific primers were used to assess the Tpm1 and Tpm2 levels in wild type and different mutant background. TDH1 and PGK1 housekeeping genes were used as controls for all qRT-PCR experiments.

### Vesicle delivery to bud

Vesicle delivery was visualized using an N-terminal tagged ymScarleti-Sec4 construct expressed under ADH promoter as an additional copy integrated into the *ura3* locus. Overnight grown yeast cells were diluted and allowed to grow until mid-log phase. Cells were adhered on a Concanavalin A-coated glass bottom dish and z-stacks were acquired with a step size of 0.5µm to cover a range of 6µm. The ratio of ymScarleti-Sec4 intensity in the bud and mother were taken and plotted to assess targeting of vesicles to the bud via the actin cable network. Images were also acquired using Andor BC43 table-top spinning disk confocal system using a 60x oil objective and scMOS camera detection. Images were deconvolved using Andor Fusion software and any adjustments for representation were done using Fiji (ImageJ)^106^.

### Mitochondrial morphology

Mitochondrial morphology was assessed with Om45-3xmCherry tagged at its genomic locus or Su9-mNeonGreen expressed from an integrated plasmid as a mitochondrial marker during imaging. Z-stacks of mid-log phase cells were acquired using Olympus IX83 widefield fluorescence microscope system and images were processed using the Mitochondria Analyzer plug-in in Fiji^106^ to quantify mitochondrial morphology. An example of the quantification procedure and details of parameters used for segmentation of mitochondria are shown in **Fig. S7B**.

### Actin patch imaging

Actin patch lifetime was measured using LifeAct-eGFP as an actin patch marker. Time-lapse images of LifeAct-eGFP expressing cells were acquired at an interval of 0.5 sec for 120 cycles with a 100x oil objective using Olympus SpinSR spinning disk confocal system. Patch lifetime was calculated as the time between appearance and disappearance of LifeAct-eGFP patch fluorescence signal. Imag

### Retrograde Actin Cable Flow

Retrograde actin cable flow was assessed using LifeAct-eGFP as an actin cable marker. Time-lapse images of LifeAct-eGFP expressing cells were acquired at an interval of 1 sec for 20 cycles with a 100x oil objective using Olympus SpinSR spinning disk confocal system. Images were analyzed to measure rate of actin cable elongation towards the rear of the mother cell.

### Latrunculin B sensitivity assay

Yeast cells were grown to mid-log phase at 25°C and LatB (SigmaAldrich, cat. No.: 428020) was added to a final concentration of 67μM. Cells were fixed with 4% paraformaldehyde at time points 0 min, 2 min and 04.30 min after LatB addition. The cells were then stained with Alexa488-phalloidin and imaged as described above.

### Image analysis and statistical analysis

All images were processed using Fiji (ImageJ)^106^. The specific analysis pipelines are mentioned in the above sections for each type of measurement. The graphs were plotted using GraphPad Prism (v.6.04) and statistical tests were also performed using in-built functions in GraphPad Prism. * p < 0.05, ** p < 0.01, *** p < 0.001, ****p < 0.0001

## Acknowledgements

We thank the Department of Biochemistry, Indian Institute of Science for access to DST-FIST imaging and other central facilities. We also thank the Divisional Bioimaging Facility. We are grateful to Mr. Pabitra Sharma for help with image analysis. We thank Prof. PN Rangarajan, Prof. Ramanujam Srinivasan, Prof. Sunil Laxman, Prof. Sachin Kotak, Prof. Sunish Radhakrishnan, and Prof. Marko Kaksonen for their feedback on the manuscript. AD and JK acknowledge GATE fellowship from IISc. JSB acknowledges KVPY fellowship from IISc. We thank Silvia Jansen for advice on quantification of TIRF microscopy experiments and Blake Miller for help with protein purification.

## Author Contributions

SP and AD conceived and supervised the experiments. AD and BVT constructed yeast strains and expression plasmids. AD and BVT performed experiments and data analysis. AS made yeast strains and performed data analysis. JK made the plasmid constructs for protein purification, performed western blotting, and data analysis. SB purified proteins and performed *in-vitro* reconstitution experiments, and data analysis. SS conceived and supervised the in-vitro experiments with purified proteins. JSB performed data analysis. SP, SS, AD and BVT wrote the manuscript, and all the authors helped in editing the manuscript.

## Funding

This work was financially supported by a Department of Biotechnology-Wellcome Trust India Alliance intermediate fellowship (IA/I/21/1/505633), SERB SRG grant (SRG/2021/001600) and an Indian Institute of Science (IISc) start-up grant awarded to S.P. S.S. is supported by NIH NIGMS grant R35GM143050.

## Competing interest statement

The authors declare no conflict of interest.

---

**Figure S1:**
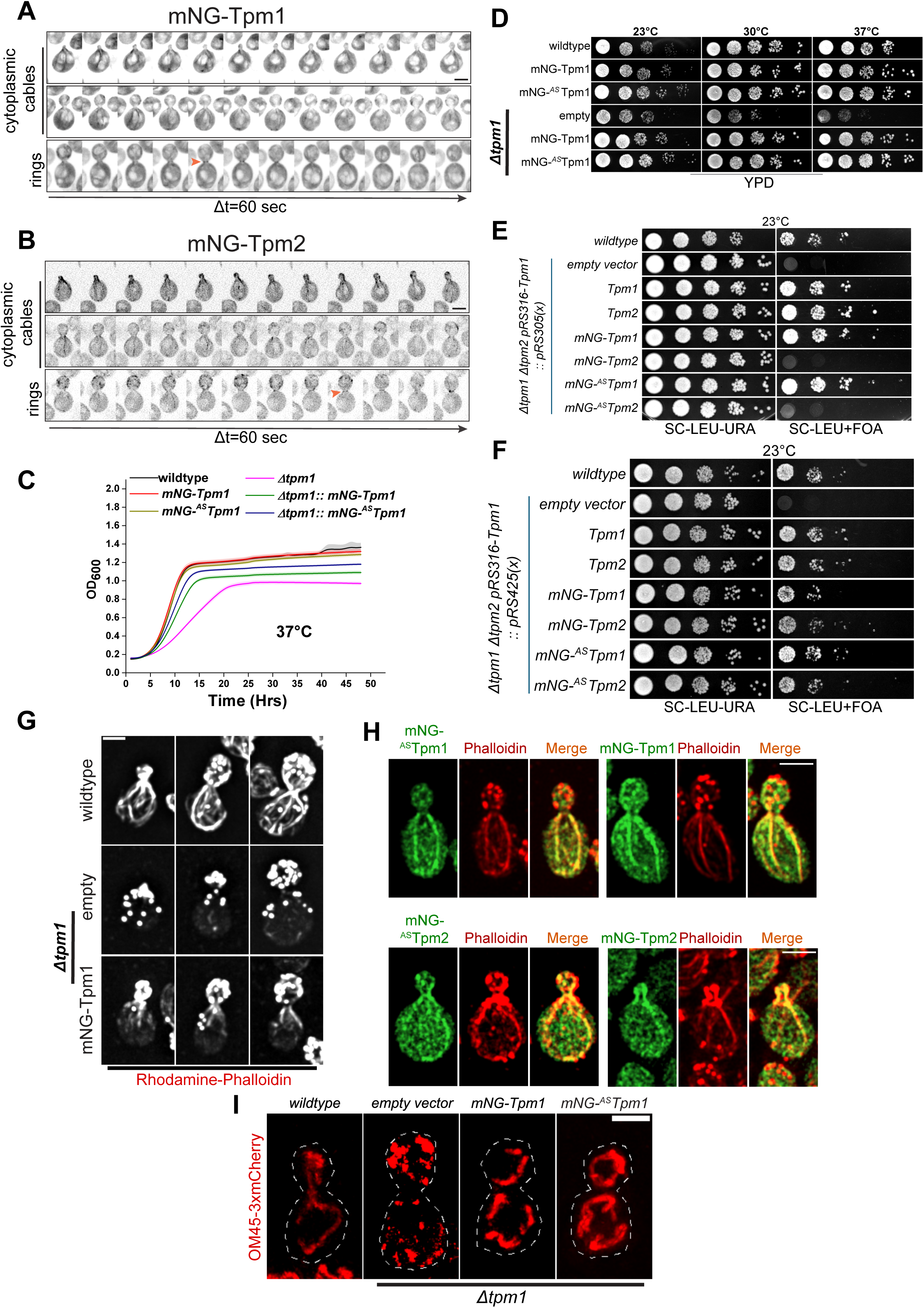
Functional characterization of mNeonGreen-Tpm fusion proteins. **(A)** Representative time-lapse montages of wildtype yeast cells expressing mNG-Tpm1; scale bar -3μm. **(B)** Representa-tive time-lapse montages of wildtype yeast cells expressing mNG-Tpm2; scale bar -3μm. **(C)** Plot representing growth curves of indicated yeast strains performed at 37°C. y-axis represents mean absorbance at 600nm. **(D)** Spot assay image for indicated yeast strains performed at 23°C, 30°C, and 37°C. **(E)** Representative images of cells of indicated yeast strains stained with Rhodamine-phalloidin; scale bar – 2μm. **(F)** Representative images of cells expressing indicated mNeonGreen-Tpm fusion proteins stained with Rhodamine-phalloidin; scale bar -2μm. **(G)** Representative images of indicated yeast strains expressing Om45-3xmCherry from the native locus; scale bar -2μm.

**Figure S2:**
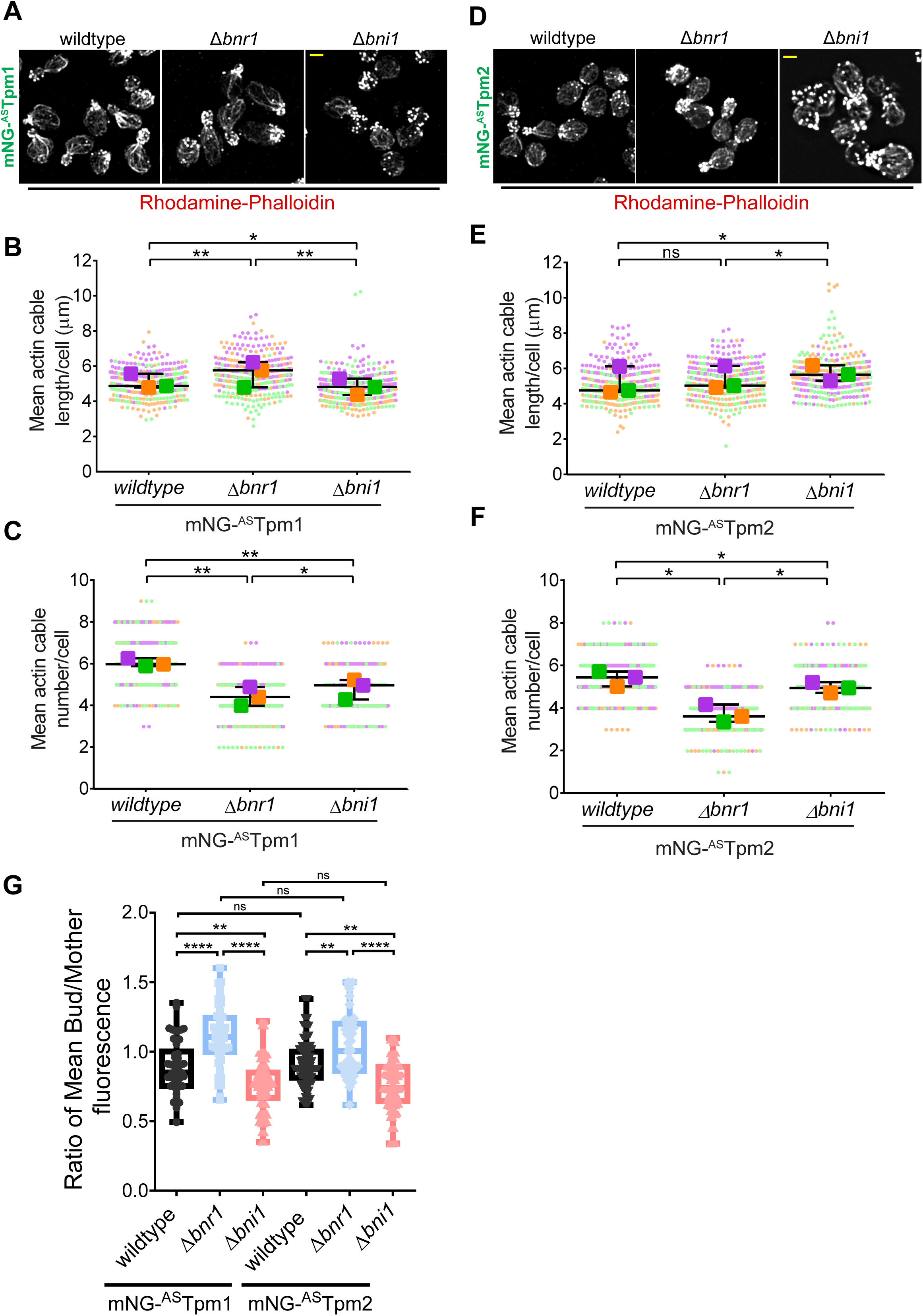
Tpm1 and Tpm2 bind to actin filaments made by formins Bnr1 and Bni1 indiscriminately. **(A)** Representative images of cells of wildtype, ∆*bnr1*, and ∆*bni1* cells expressing mNG-^AS^Tpm1 stained with Rhodamine-phalloidin; scale bar -2μm. **(B)** Superplot representing mean actin cable length per cell in wildtype, ∆*bnr1*, and ∆*bni1* cells expressing mNG-^AS^Tpm1; n=100 cells per strain per replicate, N=3. **(C)** Superplot representing mean actin cable number per cell in wildtype, ∆*bnr1*, and ∆*bni1* cells expressing mNG-^AS^Tpm1; n=100 per strain per replicate, N=3. **(D)** Representative images of cells of wildtype, ∆*bnr1*, and ∆*bni1* cells expressing mNG-^AS^Tpm2 stained with Rhodamine-phalloidin; scale bar – 2μm. **(E)** Superplot representing mean actin cable length per cell in wildtype, ∆*bnr1*, and ∆*bni1* cells expressing mNG-^AS^Tpm2; n=100 cells per strain per replicate, N=3. **(F)** Superplot representing mean actin cable number per cell in wildtype, ∆*bnr1*, and ∆*bni1* cells expressing mNG-^AS^Tpm2; n=100 cells per strain per replicate, N=3. **(G)** Box and whiskers plot showing ratio of mean mNG fluorescence in the bud to the mother compartment in the indicated strains. Box represents 25^th^ and 75^th^ percentile, line represents median, whiskers represent minimum and maximum value. (Superplots represent datapoints and means from three independent biological replicates marked in different colours; One-Way Anova with Tukey’s Multiple Comparisons test was used in (B), (C), (E), (F); * p < 0.05, ** p < 0.01)

**Figure S3:**
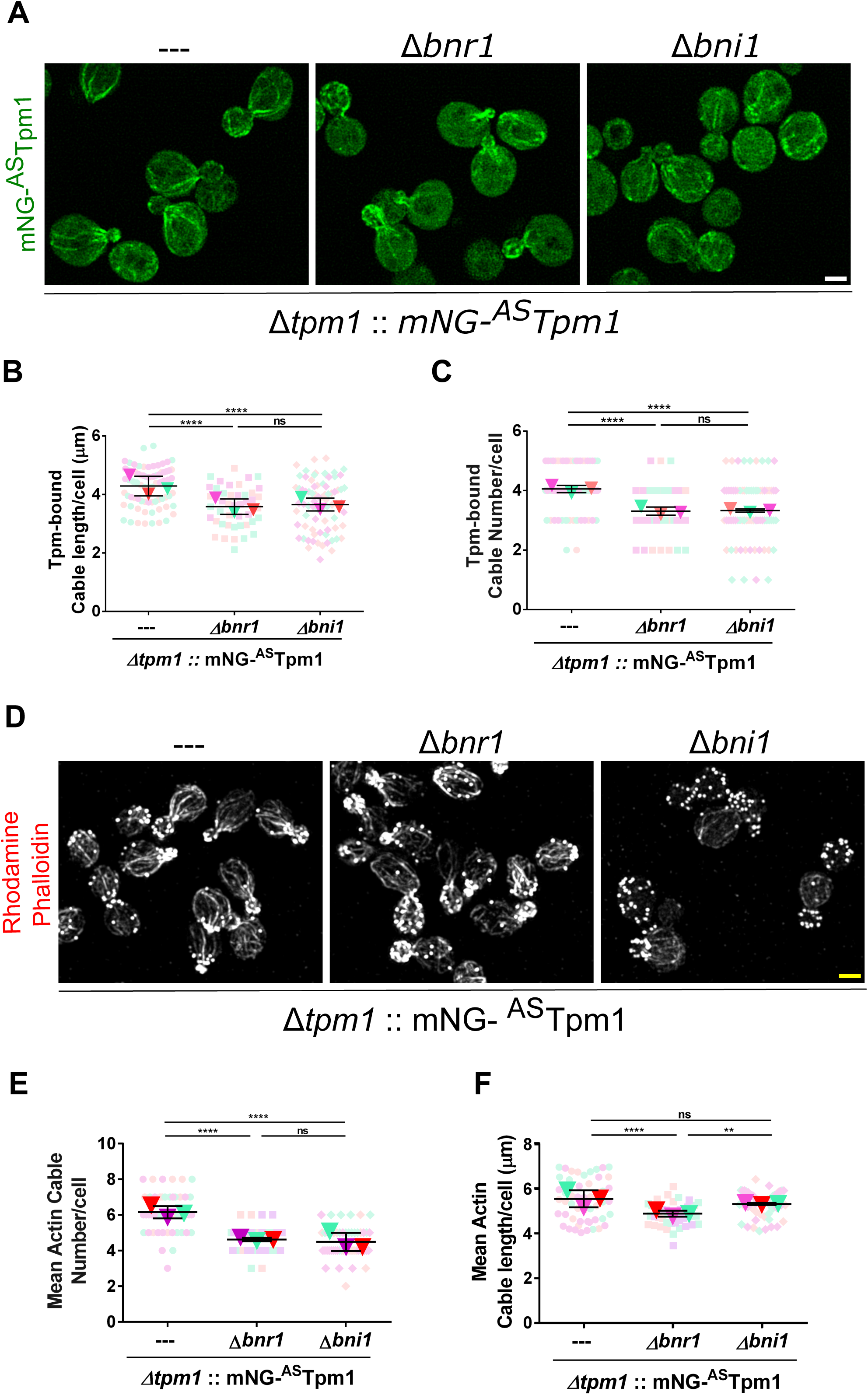
mNG-^AS^Tpm1 shows binding to both Bnr1-and Bni1-made cables even in the absence of endogenous Tpm1. **(A)** Representative images of ∆*tpm1* cells expressing mNG-^AS^Tpm1 fusion protein as sole copy with indicated genotypes; scale bar -2μm. **(B)** Superplot representing mean Tpm-bound cable length per cell in indicated strains expressing mNG-^AS^Tpm1 as sole copy; n=25 cells per strain per replicate, N=3. **(C)** Superplot representing mean Tpm-bound cable number per cell in indicated strains expressing mNG-^AS^Tpm1 as a sole copy; n=25 cells per strain per replicate, N=3. **(D)** Representative images of ∆*tpm1* cells expressing mNG-^AS^Tpm1 fusion protein as sole copy with indicated genotypes stained with Rhodamine-phalloidin; scale bar -2μm. **(E)** Superplot representing mean actin cable length per cell in indicated yeast strains shown in (D); n=15 cells per strain per replicate, N=3. **(F)** Superplot representing mean actin cable number per cell in in indicated yeast strains shown in (D); n=15 per strain per replicate, N=3. (Superplots represent datapoints and means from three independent biological replicates marked in different colours; One-Way Anova with Tukey’s Multiple Comparisons test was used in (B), (C), (E), (F); * p < 0.05, ** p < 0.01, *** p < 0.001, **** p < 0.0001)

**Figure S4:**
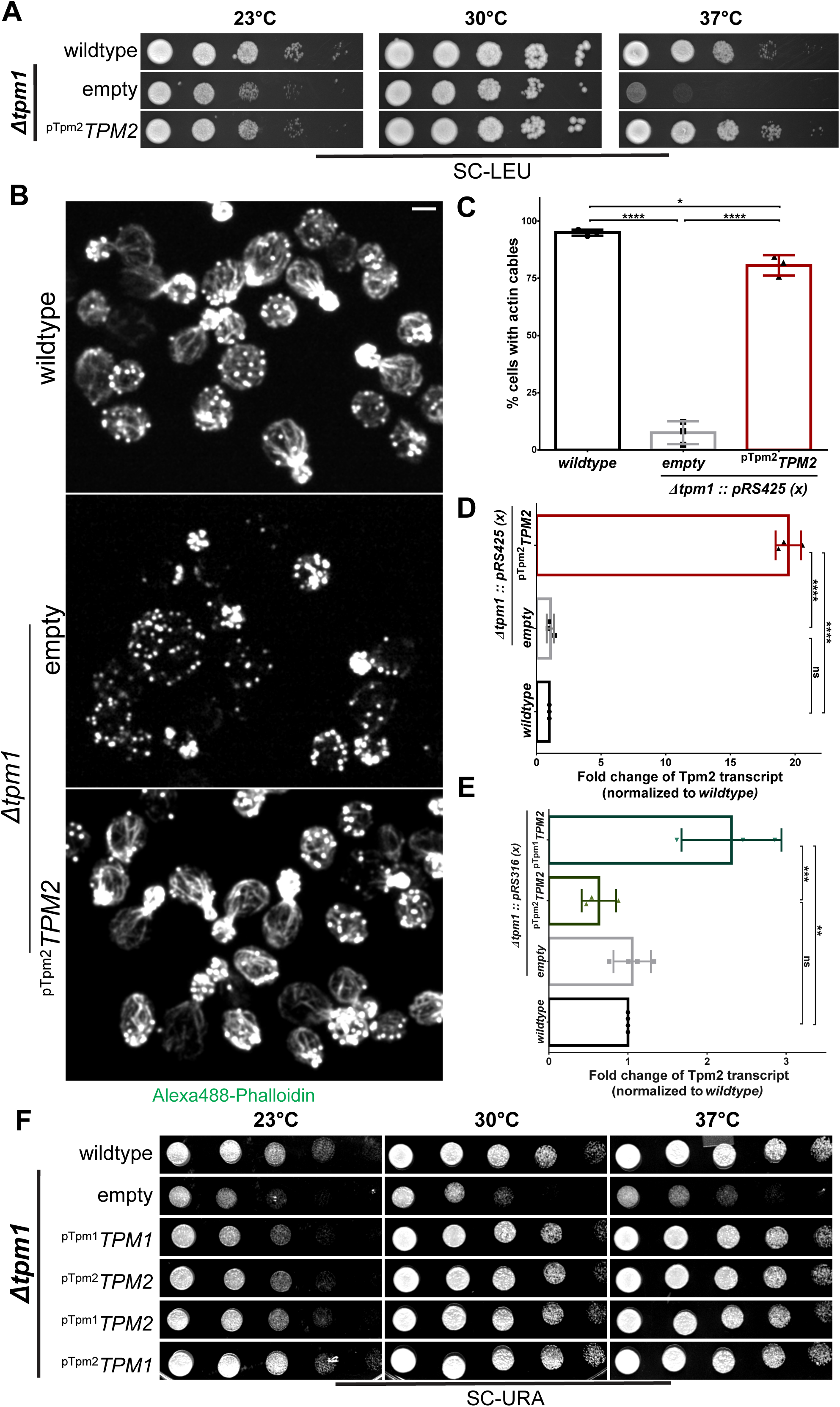
Tpm2 overexpression restores actin cables in *∆tpm1* cells. **(A)** Spot assay image for indicated yeast strains performed at 23°C, 30°C, and 37°C. **(B)** Representative images of indicated yeast strains stained with Alexa488-phalloidin; scale bar – 2μm. **(C)** Plot representing mean percentage of cells with detectable actin cables in indicated yeast strains averaged over 3 biological replicates; n≥200 cells for each strain per replicate, N=3. **(D)** Plot representing fold change of Tpm2 transcript levels normalized to wildtype in the indicated yeast strains; n=3 per strain per experiment, N=3. **(E)** Plot representing fold change of Tpm2 transcript levels normalized to *wildtype* in the indicated yeast strains; n=3 per strain per experiment, N=3. **(F)** Spot assay image for indicated yeast strains performed at 23°C, 30°C, and 37°C. (Box represents 25^th^ and 75^th^ percentile, line represents median, whiskers represent minimum and maximum value; One-Way Anova with Tukey’s Multiple Comparisons test was used in (C), (D) and (E).s * p < 0.05, ** p < 0.01, *** p < 0.001, **** p < 0.0001)

**Figure S5:**
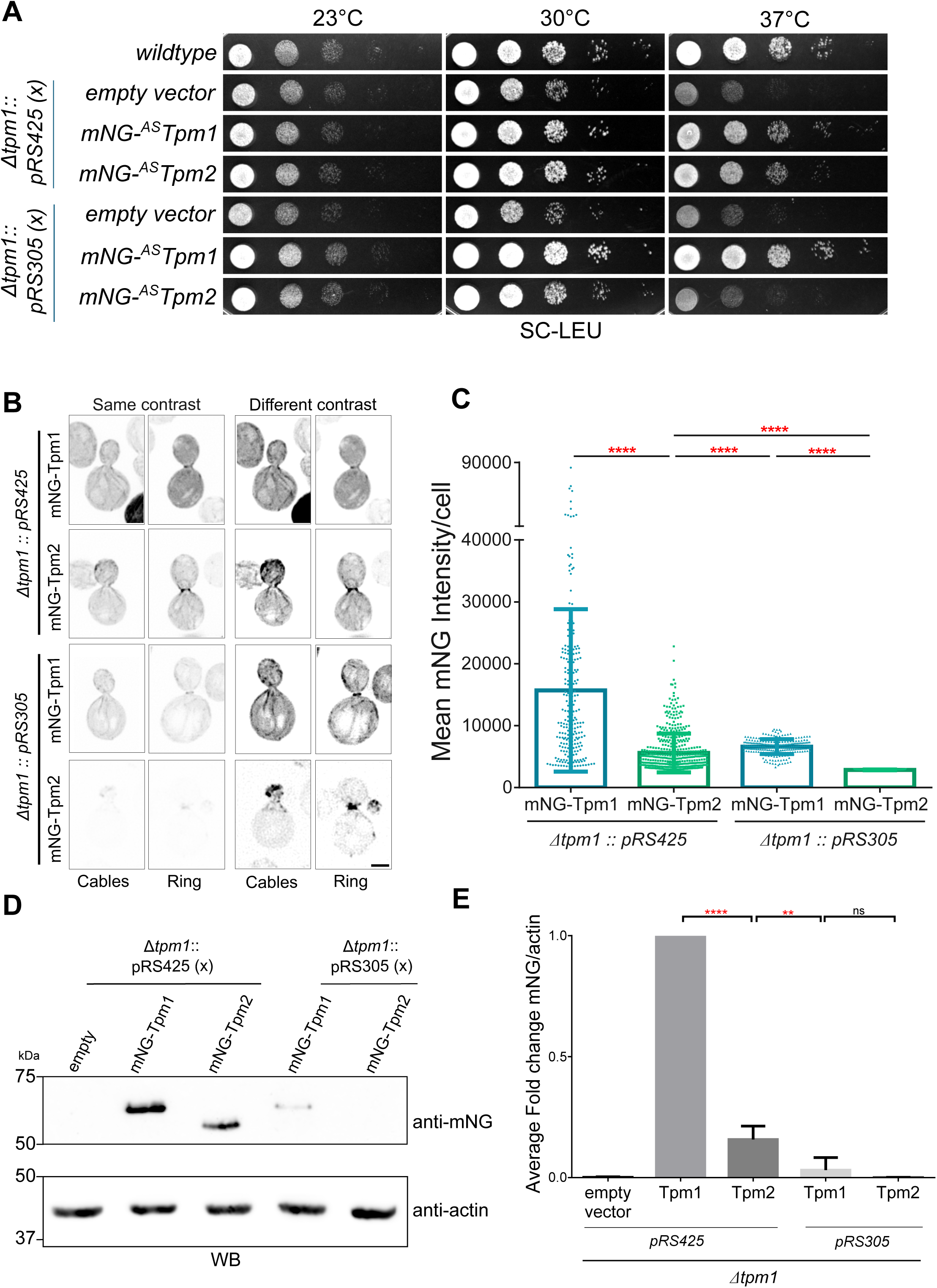
Increased protein levels of mNG-^AS^Tpm2 cause a rescue of phenotype in *∆tpm1* cells. **(A)** Spot Assay showing growth of indicated yeast strains at 23°C, 30°C, and 37°C. **(B)** Representative images of yeast cells with indicated genotypes, scale bar -2μm. **(C)** Plot showing mean mNG fluorescence intensity per cell in the indicated yeast strains, n>174 cells per strain. **(D)** Representative image of western blot probed with anti-mNG and anti-actin (loading control). **(E)** Plot showing normalized fold change of mNG/actin signal intensity for blot shown in (D), N=3. (One-Way Anova with Tukey’s Multiple Comparisons test was used in (C) and (E). * p < 0.05, ** p < 0.01, *** p < 0.001, **** p < 0.0001)

**Figure S6:**
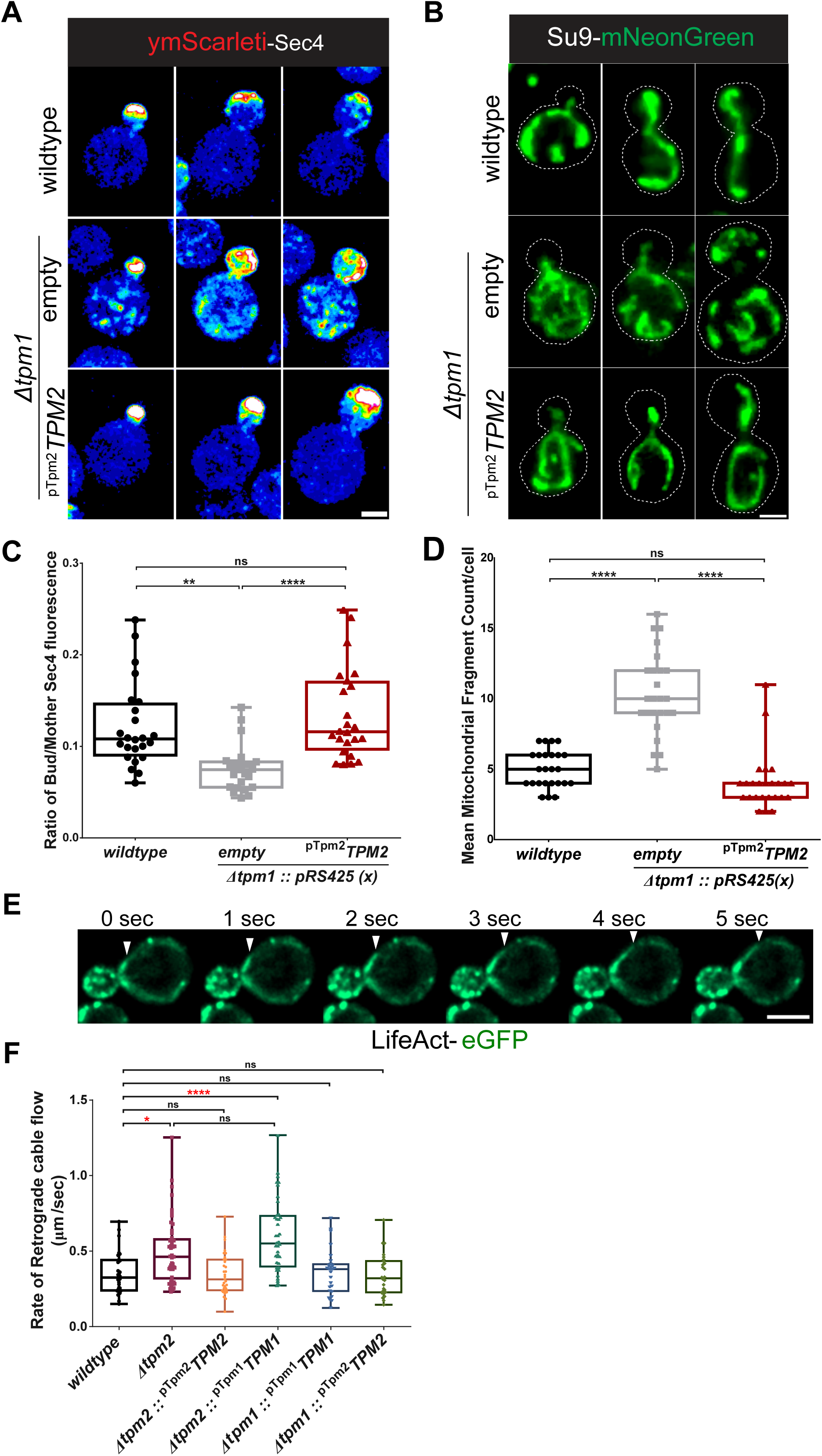
Tpm2 overexpression restores actin cable dependent functions in *∆tpm1* cells. **(A)** Representative images of indicated yeast strains expressing ymScarleti-Sec4; scale bar – 2μm. **(B)** Representative images of indicated yeast strains expressing Su9-mNeonGreen; scale bar – 2μm. **(C)** Plot representing ratio of bud/mother ymScarleti-Sec4 fluorescence per cell in the indicated yeast strains; n≥24 cells per strain. **(D)** Plot representing mean mitochondrial fragment count per cell in the indicated yeast strains; n≥25 cells per strain. **(E)** Representative time-lapse montages of wildtype yeast cells expressing LifeAct-eGFP from its native locus, scale bar -2μm. **(F)** Plot representing retrograde actin cable flow rate (μm/s) in the indicated yeast strains; n≥20 events per strain. (Box represents 25^th^ and 75^th^ percentile, line represents median, whiskers represent minimum and maximum value; One-Way Anova with Tukey’s Multiple Comparisons test was used in (C), (F); Krus-kal-Wallis test with Dunn’s multiple comparisons was used in (D); * p < 0.05, ** p < 0.01, *** p < 0.001, **** p < 0.0001)

**Figure S7:**
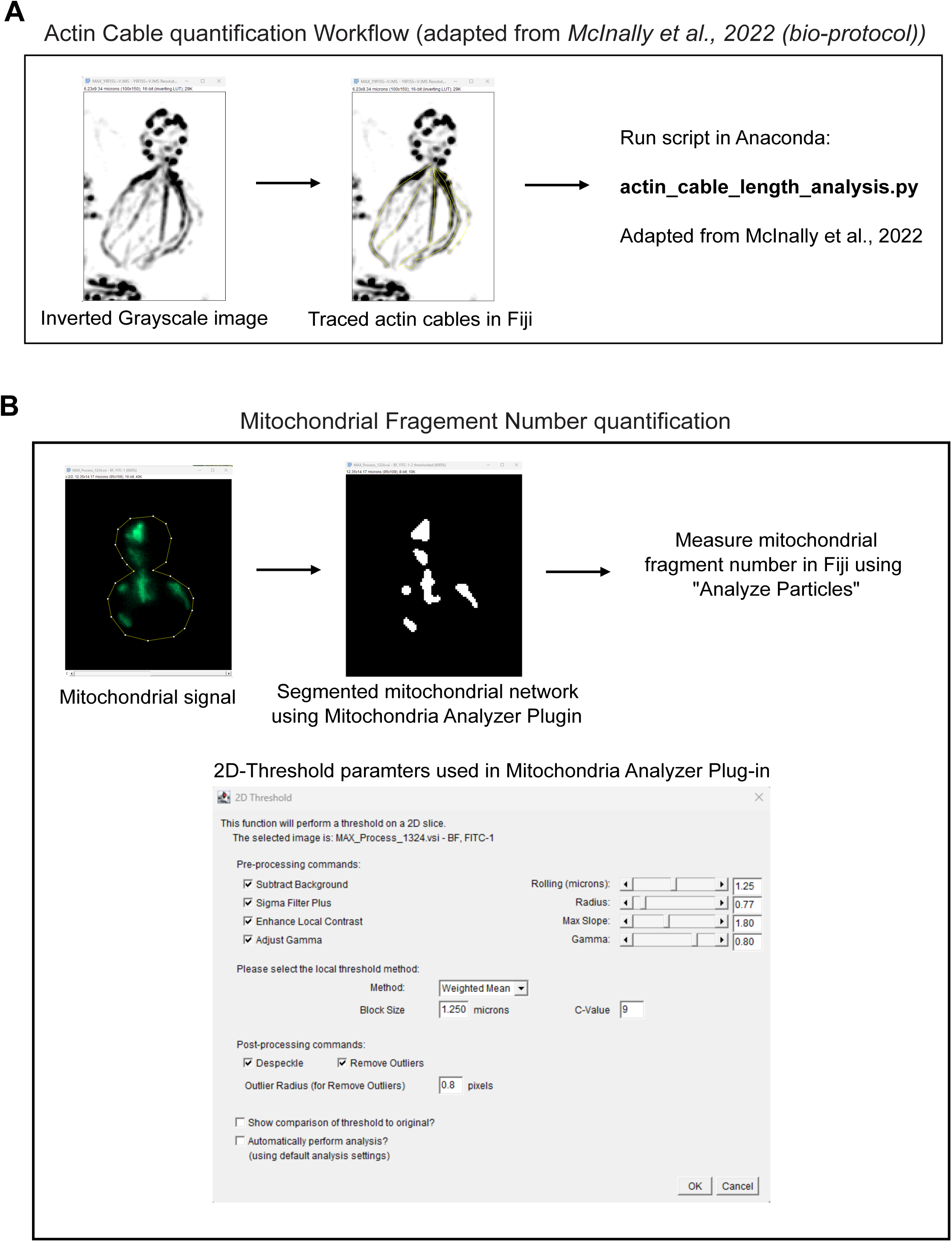
Analysis piplelines used for quantitative analysis of actin cables and mitochondrial morphology in our study. **(A)** Schematic showing quantification workflow for actin cables as adapted from *McInally et al. 2022* (*bio-protocol*). **(B)** Schematic showing quantification workflow for mitochondrial morphology analysis using Mitochondria Analyzer Plug-In in Fiji.

**Supplementary Movie 1**. Time-lapse imaging of small-, medium-and large-budded cells expressing mNeonGreen-^AS^Tpm1. (Scale bar -3μm)

**Supplementary Movie 2**. Time-lapse imaging of small-, medium-and large-budded cells expressing mNeonGreen-^AS^Tpm2. (Scale bar -3μm)

**Supplementary Movie 2**. Time-lapse imaging of haploid yeast cell expressing mNeonGreen-^AS^Tpm1 mating with another wildtype haploid cell. (Scale bar -3μm)

**Supplementary Movie 2**. Time-lapse imaging of haploid yeast cell expressing mNeonGreen-^AS^Tpm2 mating with another wildtype haploid cell. (Scale bar -3μm)

**Supplementary Table 1.** List of plasmids used in this study.

| Plasmid Number | Description | Source |
| --- | --- | --- |
| piSP1 | *pRS305* | *Sikorski and Hieter, 1989*<sup>102</sup> |
| piSP5 | *pRS315* | *Sikorski and Hieter, 1989*<sup>102</sup> |
| piSP6 | *pRS316* | *Sikorski and Hieter, 1989*<sup>102</sup> |
| piSP11 | *pRS425* | *Christianson et al. 1992*<sup>107</sup> |
| piSP14 | *pYM25 (yeGFP-hphNT2)* | *Janke et al. 2004*<sup>108</sup> |
| piSP346 | *pRS305-pTpm1-mNG-40L-Tpm1-tCYC* | This Study |
| piSP348 | *pRS305-pTpm2-mNG-40L-Tpm2-tCYC* | This Study |
| piSP1422 | *pRS305-pTpm1-mNG-40L-<sup>AS</sup>Tpm1-tTpm1* | This Study |
| piSP1423 | *pRS305-pTpm2-mNG-40L-<sup>AS</sup>Tpm2-tTpm2* | This Study |
| piSP1430 | *pRS316-pTpm1-TPM1-tTpm1* | This Study |
| piSP1493 | *pRS425-pTpm2-TPM2-tTpm2* | This Study |
| piSP1450 | *pRS316-pTpm1-TPM2-tTpm2* | This Study |
| piSP1451 | *pRS306-pADH-ymScarlet-Sec4* | This Study |
| piSP1529 | *pRS316-pTpm2-TPM2-tTpm2* | This Study |
| piSP1534 | *pET28a-6His-Cof1* | This Study |
| piSP1541 | *pRS316-pTpm2-TPM1-tTpm1* | This Study |
| piSP1644 | *pETMCN-<sup>AS</sup>Tpm1* | This Study |
| piSP1646 | *pETMCN-<sup>AS</sup>Tpm2* | This Study |
| piSP1825 | *pRS305-pADH-ymScarlet-Sec4-tCYC* | This Study |
| piSP1826 | *pRS315-pTpm1-TPM2-tTpm2* | This Study |
| piSP1827 | *pRS315-pTpm2-TPM1-tTpm1* | This Study |
| piSP1828 | *pRS315-pTpm2-TPM2-tTpm2* | This Study |
| piSP1829 | *pRS305-pCYC-Su9-mNG-tCYC* | This Study |
| piSP1830 | *pRS306-pCYC-Su9-mNG-tCYC* | This Study |
| piSP1831 | *pRS315-pTpm1-TPM1-tTpm1* | This Study |
| piSP1836 | *pRS306-pTpm1-ymScarleti-40L-<sup>AS</sup>Tpm1-tTpm1* | This Study |
| piSP2034 | *pRS305-pADH-LifeAct-eGFP-tCYC* | This Study |

**Supplementary Table 2.** List of yeast strains used in this study.

| Strain Number | Genotype | Source |
| --- | --- | --- |
| YSP002 | *MATa ura3-52 leu2Δ1 trp1Δ63 his3Δ200* | *Pereira and Schiebel, 2001*<sup>109</sup> |
| YSP003 | *MATα ura3-52 lys2-801amber ade2-101ochre trp1Δ63 his3Δ200 leu2Δ1* | *Sikorski and Hieter, 1989*<sup>102</sup> |
| YSP107 | *MATa ura3-52 leu2Δ1 trp1Δ63 his3Δ200 pRS305-pTpm1-mNG-40aaL-Tpm1-tTpm1* | This study |
| YSP108 | *MATa ura3-52 leu2Δ1 trp1Δ63 his3Δ200 pRS305-pTpm2-mNG-40aaL-Tpm2-tTpm2* | This study |
| YSP146 | *MATa ura3-52 leu2Δ1 trp1Δ63 his3Δ200 -Δtpm1-HIS3MX6-pRS305-pTpm1-mNG-40L-Tpm1-tTpm1* | This study |
| YSP195 | *MATa ura3-52 leu2Δ1 trp1Δ63 his3Δ200-Δtpm1-HIS3MX6* |  |
| YSP196 | *MATa ura3-52 leu2Δ1 trp1Δ63 his3Δ200-Δtpm2-HIS3MX6* |  |
| YSP227 | *MATa ura3-52 leu2Δ1 trp1Δ63 his3Δ200 -Δtpm1-HIS3MX6-pRS305* | This study |
| YSP355 | *MATa ura3-52 leu2Δ1 trp1Δ63 his3Δ200 -pRS305-pTpm1-mNG-40aaL-(AM)Tpm1-tTpm1* | This study |
| YSP356 | *MATa ura3-52 leu2Δ1 trp1Δ63 his3Δ200 -pRS305-pTpm2-mNG-40aaL-(am)Tpm2-tTpm2* | This study |
| YSP419 | *MATa ura3-52 leu2Δ1 trp1Δ63 his3Δ200 -Δtpm1-HIS3MX6-pRS305-pTpm1-mNG-40L-<sup>AS</sup>Tpm1-tTpm1* | This study |
| YSP440 | *MATa ura3-52 leu2Δ1 trp1Δ63 his3Δ200 -pRS305-pTpm1-mNG-40aaL-(am)Tpm1-tTpm1 -Δbnr1-HIS3MX6* | This study |
| YSP441 | *MATa ura3-52 leu2Δ1 trp1Δ63 his3Δ200 -pRS305-pTpm2-mNG-40aaL-(am)Tpm2-tTpm2-Δbnr1-HIS3MX6* | This study |
| YSP442 | *MATa ura3-52 leu2Δ1 trp1Δ63 his3Δ200 -pRS305-pTpm2-mNG-40aaL-Tpm2-tTp-Δbni1-HIS3MX6* | This study |
| YSP443 | *MATa ura3-52 leu2Δ1 trp1Δ63 his3Δ200 -pRS305-pTpm1-mNG-40aaL-(am)Tpm1-tTpm1 -Δbni1-HIS3MX6* | This study |
| YSP609 | *MATa ura3-52 leu2Δ1 trp1Δ63 his3Δ200 -Δtpm1-HIS3MX6-pRS305-pTpm1-mNG-40L-Tpm1-tTpm1 - OM45-3XmCherry-Kan* | This study |
| YSP610 | *MATa ura3-52 leu2Δ1 trp1Δ63 his3Δ200 -Δtpm1-HIS3MX6-OM45-3XmCherry-Kan* | This study |
| YSP611 | *MATa ura3-52 leu2Δ1 trp1Δ63 his3Δ200 -Δtpm1-HIS3MX6-pRS305-pTpm1-mNG-40L-<sup>AS</sup>Tpm1-tTpm1 -OM45-3XmCherry-Kan* | This study |
| YSP612 | *MATa ura3-52 leu2Δ1 trp1Δ63 his3Δ200 --OM45-3XmCherry-Kan* | This study |
| YSP671 | *MATa ura3-52 leu2Δ1 trp1Δ63 his3Δ200 -pRS305-pTpm2-mNG-40aaL-(AM)Tpm2-tTpm2 - pRS306-pTpm1-ymScarletil-40aaLinker-(AM)Tpm1-tTpm1* | This study |
| YSP685 | *MATa ura3-52 leu2Δ1 trp1Δ63 his3Δ200 - Δtpm1-HIS3MX6-pRS425* | This study |
| YSP686 | *MATa ura3-52 leu2Δ1 trp1Δ63 his3Δ200 -Δtpm1-HIS3MX6-pRS425-pTpm2-Tpm2-tTpm2* | This study |
| YSP767 | *MATa ura3-52 leu2Δ1 trp1Δ63 his3Δ200 -Δtpm1-HIS3MX6 - pRS425-pTpm2-mNG-40L-<sup>AS</sup>Tpm2-tTpm2* | This study |
| YSP780 | *MATa ura3-52 leu2Δ1 trp1Δ63 his3Δ200 -Δtpm1-his3MX6-pRS305-pTpm1-mNG-40L-<sup>AS</sup>Tpm1-tTpm1 - Δbnr1-hphNT1* | This study |
| YSP794 | *MATa ura3-52 leu2Δ1 trp1Δ63 his3Δ200 -Δtpm1-HIS3MX6 - pRS425-pTpm1-mNG-40L-<sup>AS</sup>Tpm1-tTpm1* | This study |
| YSP824 | *MATa ura3-52 leu2Δ1 trp1Δ63 his3Δ200 -Δtpm1-HIS3MX6 - pRS306-pADH-ymScarleti-Sec4* | This study |
| YSP835 | *MATa ura3-52 leu2Δ1 trp1Δ63 his3Δ200 - pRS425 - pRS316* | This study |
| YSP836 | *MATa ura3-52 leu2Δ1 trp1Δ63 his3Δ200 - pRS306-pADH-ymScarleti-Sec4* | This study |
| YSP837 | *MATa ura3-52 leu2Δ1 trp1Δ63 his3Δ200 -Δtpm1-HIS3MX6-pRS425-pTpm2-Tpm2-tTpm2 - pRS306-pADH-ymScarleti-Sec4* | This study |
| YSP848 | *MATa ura3-52 leu2Δ1 trp1Δ63 his3Δ200 -Δtpm1-HIS3MX6 - pRS316-pTpm2-Tpm1-tTpm1* | This study |
| YSP887 | *MATa ura3-52 leu2Δ1 trp1Δ63 his3Δ200 -Δtpm1-HIS3MX6 - pRS316* | This study |
| YSP907 | *MATa ura3-52 leu2Δ1 trp1Δ63 his3Δ200Δtpm1-HIS3MX6; pRS316-pTpm1-Tpm1-tTpm1* | This study |
| YSP909 | *MATa ura3-52 leu2Δ1 trp1Δ63 his3Δ200Δtpm1-HIS3MX6; pRS316-pTpm2-Tpm2-tTpm2* | This study |
| YSP1090 | *MATa ura3-52 leu2Δ1 trp1Δ63 his3Δ200-Δtpm1-HIS3MX6; pRS316-pTpm1-Tpm2-tTpm2* | This study |
| YSP1117 | *MATa ura3-52 leu2Δ1 trp1Δ63 his3Δ200-pRS306-pCYC-Su9-mNG-tCYC - pRS425* | This study |
| YSP1118 | *MATa ura3-52 leu2Δ1 trp1Δ63 his3Δ200-Δtpm1-HIS3MX6-pRS425 - pRS306-pCYC-Su9-mNG-tCYC* | This study |
| YSP1119 | *MATa ura3-52 leu2Δ1 trp1Δ63 his3Δ200-Δtpm1-HIS3MX6-pRS425-pTpm2-Tpm2-tTpm2 - pRS306-pCYC-Su9-mNG-tCYC* | This study |
| YSP1120 | *MATa ura3-52 leu2Δ1 trp1Δ63 his3Δ200-Δtpm1-HIS3MX6 - pRS306-pADH-ymScarleti-Sec4 - pRS315* | This study |
| YSP1121 | *MATa ura3-52 leu2Δ1 trp1Δ63 his3Δ200-Δtpm1-HIS3MX6 - pRS306-pADH-ymScarleti-Sec4 - pRS315-pTpm1-Tpm2-tTpm2* | This study |
| YSP1122 | *MATa ura3-52 leu2Δ1 trp1Δ63 his3Δ200-Δtpm1-HIS3MX6 - pRS306-pADH-ymScarleti-Sec4 - pRS315-pTpm2-Tpm1-tTpm1* | This study |
| YSP1123 | *MATa ura3-52 leu2Δ1 trp1Δ63 his3Δ200-Δtpm1-HIS3MX6 - pRS306-pADH-ymScarleti-Sec4 - pRS315-pTpm2-Tpm2-tTpm2* | This study |
| YSP1126 | *MATa ura3-52 leu2Δ1 trp1Δ63 his3Δ200 pRS305-pCYC-Su9-mNG-tCYC pRS216* | This study |
| YSP1127 | *MATa ura3-52 leu2Δ1 trp1Δ63 his3Δ200 Δtpm1-HIS3MX6 - pRS305-pCYC-Su9-mNG-tCYC* | This study |
| YSP1128 | *MATa ura3-52 leu2Δ1 trp1Δ63 his3Δ200 Δtpm1-HIS3MX6 - pRS316-pTpm2-Tpm1-tTpm1 - pRS305-pCYC-Su9-mNG-tCYC* | This study |
| YSP1129 | *MATa ura3-52 leu2Δ1 trp1Δ63 his3Δ200 Δtpm1-HIS3MX6; pRS316-pTpm1-Tpm1-tTpm1- pRS305-pCYC-Su9-mNG-tCYC* | This study |
| YSP1130 | *MATa ura3-52 leu2Δ1 trp1Δ63 his3Δ200 Δtpm1-HIS3MX6; pRS316-pTpm2-Tpm2-tTpm2- pRS305-pCYC-Su9-mNG-tCYC* | This study |
| YSP1131 | *MATa ura3-52 leu2Δ1 trp1Δ63 his3Δ200 Δtpm1-HIS3MX6; pRS316-pTpm1-Tpm2-tTpm2- pRS305-pCYC-Su9-mNG-tCYC* | This study |
| YSP1150 | *MATa ura3-52 leu2Δ1 trp1Δ63 his3Δ200-Δtpm1-HIS3MX6 - pRS306-pADH-ymScarleti-Sec4 , pRS315-pTpm1-Tpm1-tTpm1* | This study |
| YSP1189 | *MATa ura3-52 leu2Δ1 trp1Δ63 his3Δ200-Abp140-GFP-KanMX6* | This study |
| YSP1191 | *MATa ura3-52 leu2Δ1 trp1Δ63 his3Δ200-Δtpm2-HIS3MX6 - Abp140-GFP-KanMX6* | This study |
| YSP1192 | *MATa ura3-52 leu2Δ1 trp1Δ63 his3Δ200-Δtpm2-HIS3MX6) - pRS425-pTpm2-Tpm2-tTpm2) - Abp140-GFP-KanMX6* | This study |
| YSP1577 | *MATa ura3-52 leu2Δ1 trp1Δ63 his3Δ200 Δtpm1-His3MX6-Δbni1-hph - pRS305-pTpm1-mNG-40L<sup>AS</sup>Tpm1-tTpm* | This study |
| YSP1639 | *MATa ura3-52 leu2Δ1 trp1Δ63 his3Δ200 - Δtpm1-his3MX6 - pRS305-pTpm2-mNG<sup>AS</sup>Tpm2-tTpm2* | This study |
| YSP1816 | *MATa ura3-52 leu2Δ1 trp1Δ63 his3Δ200 pRS305-pADH-Life-Act-eGFP-tCYC - Δtpm1::his3MX6 - pRS316 (empty)* | This study |
| YSP1817 | *MATa ura3-52 leu2Δ1 trp1Δ63 his3Δ200 pRS305-pADH-Life-Act-eGFP-tCYC - Δtpm2::his3MX6 - pRS316 (empty)* | This study |
| YSP1818 | *MATa ura3-52 leu2Δ1 trp1Δ63 his3Δ200 pRS305-pADH-Life-Act-eGFP-tCYC - pRS316 (empty)* | This study |
| YSP1819 | *MATa ura3-52 leu2Δ1 trp1Δ63 his3Δ200 pRS305-pADH-Life-Act-eGFP-tCYC - Δtpm1::his3MX6 - pRS316-pTpm1-Tpm1-tTpm1 (GD13)* | This study |
| YSP1820 | *MATa ura3-52 leu2Δ1 trp1Δ63 his3Δ200 pRS305-pADH-Life-Act-eGFP-tCYC - Δtpm2::his3MX6 - pRS316-pTpm1-Tpm1-tTpm1 (GD13)* | This study |
| YSP1822 | *MATa ura3-52 leu2Δ1 trp1Δ63 his3Δ200 pRS305-pADH-Life-Act-eGFP-tCYC - Δtpm1::his3MX6 - pRS316-pTpm1-Tpm1-tTpm1 (GT34)* | This study |
| YSP1823 | *MATa ura3-52 leu2Δ1 trp1Δ63 his3Δ200 pRS305-pADH-Life-Act-eGFP-tCYC - Δtpm2::his3MX6 - pRS316-pTpm1-Tpm1-tTpm1 (GT34)* | This study |

## Notes

### Competing Interest Statement

The authors have declared no competing interest.

### Summary of Updates

The revised manuscript throughly addressed the reviewers comments from ReviewCommons. Detailed response of the changes can be found in the authors response to reviewers file/section.

