## Supplemental figures for "Functional redundancy and formin-independent localization of tropomyosin isoforms in Saccharomyces cerevisiae"

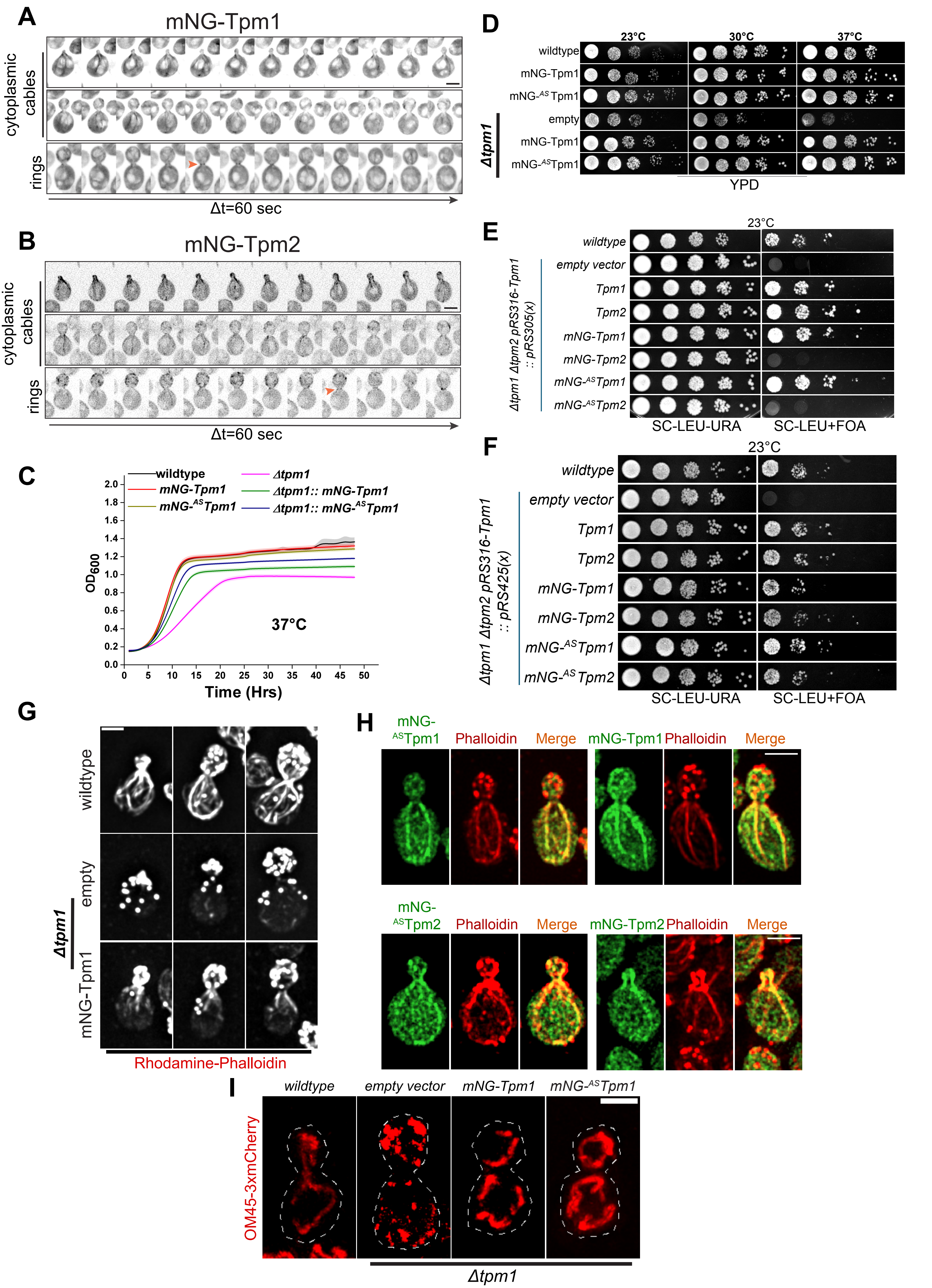
A
mNG-Tpm1
cytoplasmic cables
rings
Δt=60 sec
B
mNG-Tpm2
cytoplasmic cables
rings
Δt=60 sec
C
wildtype
mNG-Tpm1
mNG-^AS Tpm1
Δtpm1
Δtpm1:: mNG-Tpm1
Δtpm1:: mNG-^AS Tpm1
OD600
37°C
Time (Hrs)
D
23°C
30°C
37°C
wildtype
mNG-Tpm1
mNG-^AS Tpm1
empty
mNG-Tpm1
mNG-^AS Tpm1
Δtpm1
YPD
E
23°C
wildtype
empty vector
Tpm1
Tpm2
mNG-Tpm1
mNG-Tpm2
mNG-^AS Tpm1
mNG-^AS Tpm2
Δtpm1 Δtpm2 pRS316-Tpm1 :: pRS305(x)
SC-LEU-URA
SC-LEU+FOA
F
23°C
wildtype
empty vector
Tpm1
Tpm2
mNG-Tpm1
mNG-Tpm2
mNG-^AS Tpm1
mNG-^AS Tpm2
Δtpm1 Δtpm2 pRS316-Tpm1 :: pRS425(x)
SC-LEU-URA
SC-LEU+FOA
G
wildtype
empty
mNG-Tpm1
Δtpm1
Rhodamine-Phalloidin
H
mNG-^AS Tpm1
Phalloidin
Merge
mNG-Tpm1
Phalloidin
Merge
mNG-^AS Tpm2
Phalloidin
Merge
mNG-Tpm2
Phalloidin
Merge
I
wildtype
empty vector
mNG-Tpm1
mNG-^AS Tpm1
OM45-3xmCherry
Δtpm1

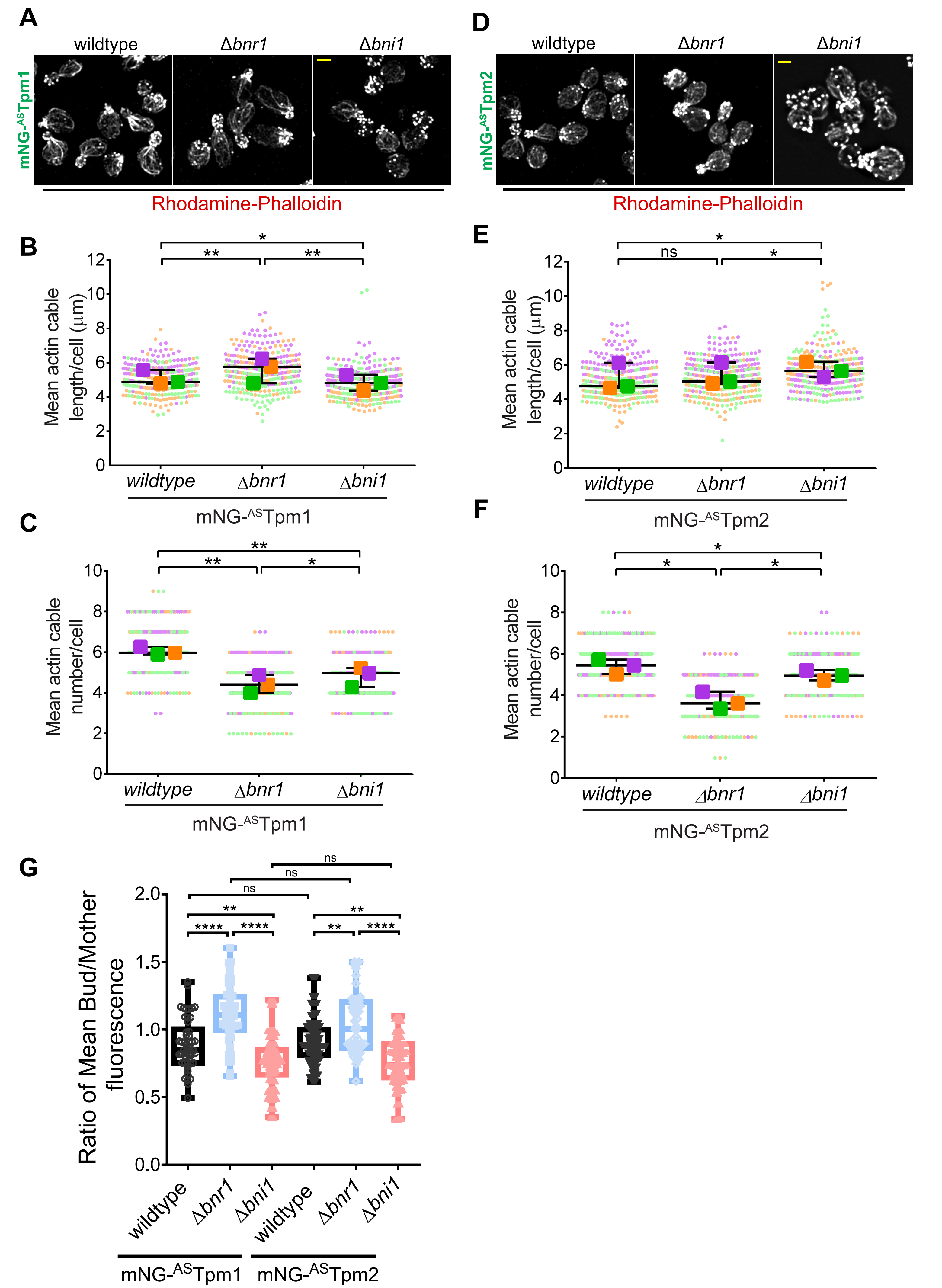
A
wildtype
Δbnr1
Δbni1
mNG-ASTpm1
Rhodamine-Phalloidin
D
wildtype
Δbnr1
Δbni1
mNG-ASTpm2
Rhodamine-Phalloidin
B
Mean actin cable length/cell (μm)
wildtype
Δbnr1
Δbni1
mNG-ASTpm1
E
Mean actin cable length/cell (μm)
ns
wildtype
Δbnr1
Δbni1
mNG-ASTpm2
C
Mean actin cable number/cell
wildtype
Δbnr1
Δbni1
mNG-ASTpm1
F
Mean actin cable number/cell
wildtype
Δbnr1
Δbni1
mNG-ASTpm2
G
Ratio of Mean Bud/Mother fluorescence
ns
ns
ns
**
****
****
**
**
****
wildtype
Δbnr1
Δbni1
wildtype
Δbnr1
Δbni1
mNG-ASTpm1
mNG-ASTpm2

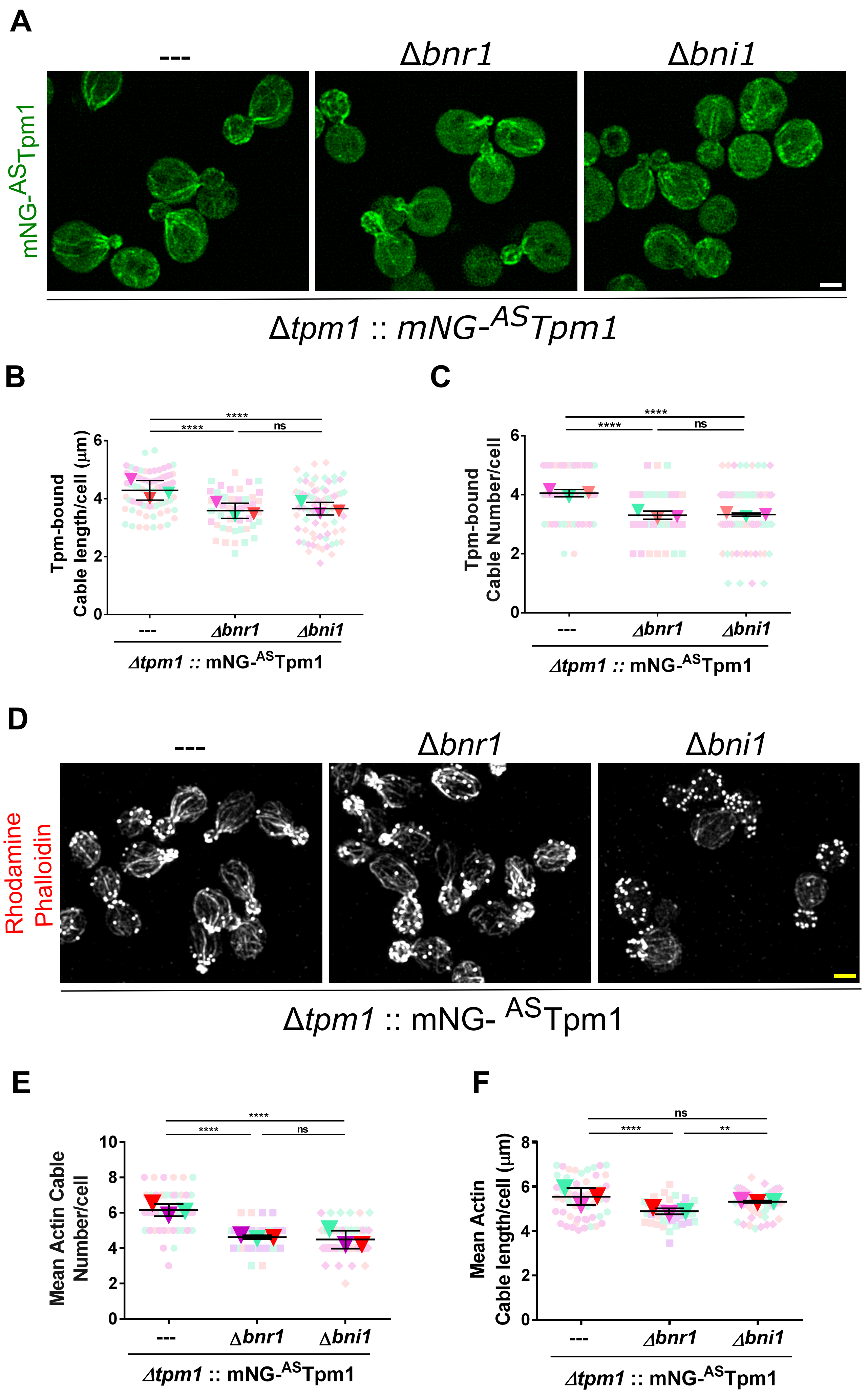
A
---
Δbnr1
Δbni1
mNG-ASTpm1
Δtpm1 :: mNG-ASTpm1
B
Tpm-bound Cable length/cell (μm)
****
****
ns
---
Δbnr1
Δbni1
Δtpm1 :: mNG-ASTpm1
C
Tpm-bound Cable Number/cell
****
****
ns
---
Δbnr1
Δbni1
Δtpm1 :: mNG-ASTpm1
D
---
Δbnr1
Δbni1
Rhodamine Phalloidin
Δtpm1 :: mNG- ASTpm1
E
Mean Actin Cable Number/cell
****
****
ns
---
Δbnr1
Δbni1
Δtpm1 :: mNG-ASTpm1
F
Mean Actin Cable length/cell (μm)
ns
****
**
---
Δbnr1
Δbni1
Δtpm1 :: mNG-ASTpm1

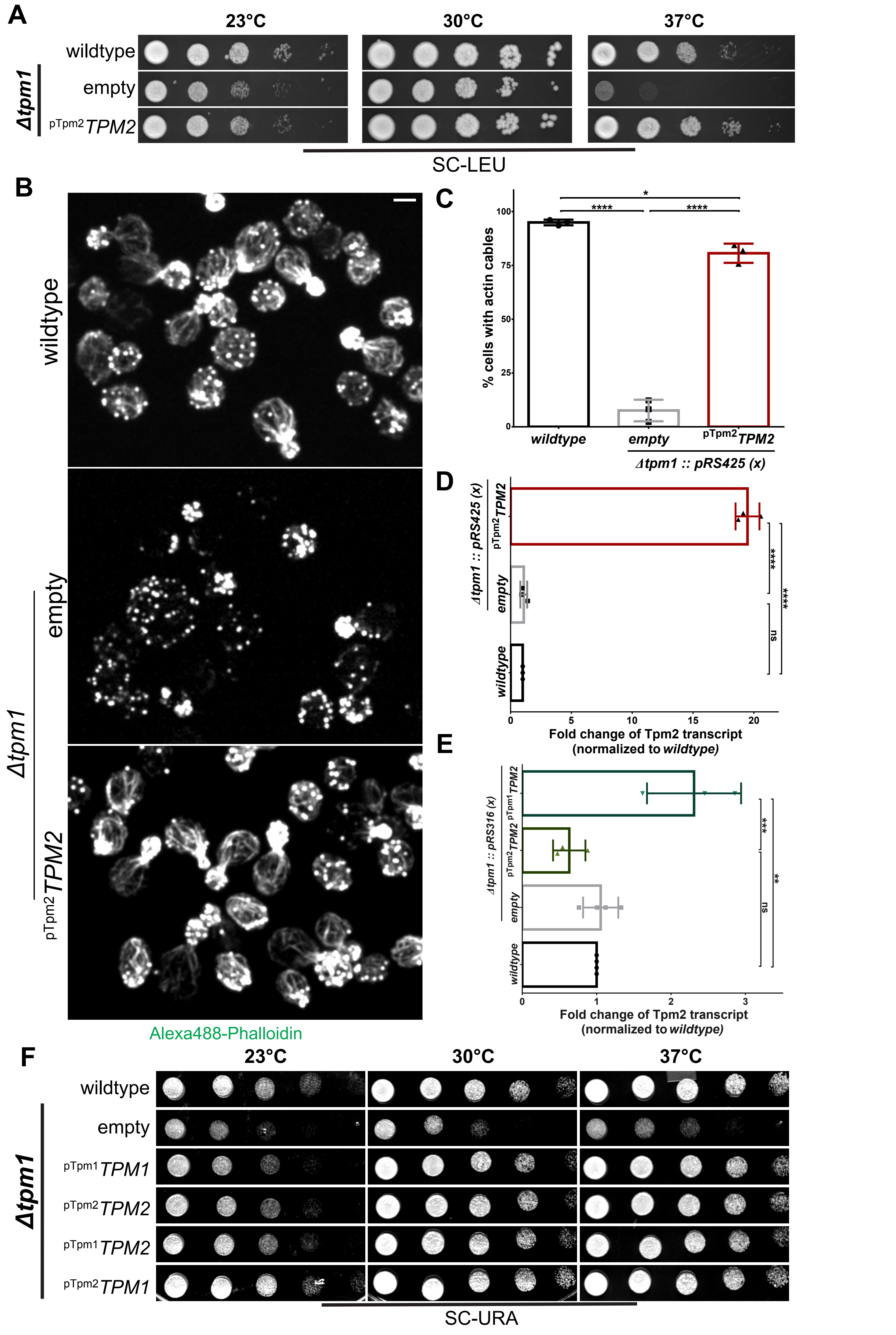

A
23°C
30°C
37°C
wildtype
Δtpm1
empty
pTpm2 TPM2
SC-LEU
B
wildtype
Δtpm1
empty
pTpm2 TPM2
Alexa488-Phalloidin
C
% cells with actin cables
100
75
50
25
0
*
****
****
wildtype
empty
pTpm2 TPM2
Δtpm1 :: pRS425 (x)
D
Δtpm1 :: pRS425 (x)
pTpm2 TPM2
empty
wildtype
****
****
ns
0
5
10
15
20
Fold change of Tpm2 transcript
(normalized to wildtype)
E
Δtpm1 :: pRS316 (x)
pTpm1 TPM2
pTpm2 TPM2
empty
wildtype
***
**
ns
0
1
2
3
Fold change of Tpm2 transcript
(normalized to wildtype)
F
23°C
30°C
37°C
wildtype
Δtpm1
empty
pTpm1 TPM1
pTpm2 TPM2
pTpm1 TPM2
pTpm2 TPM1
SC-URA

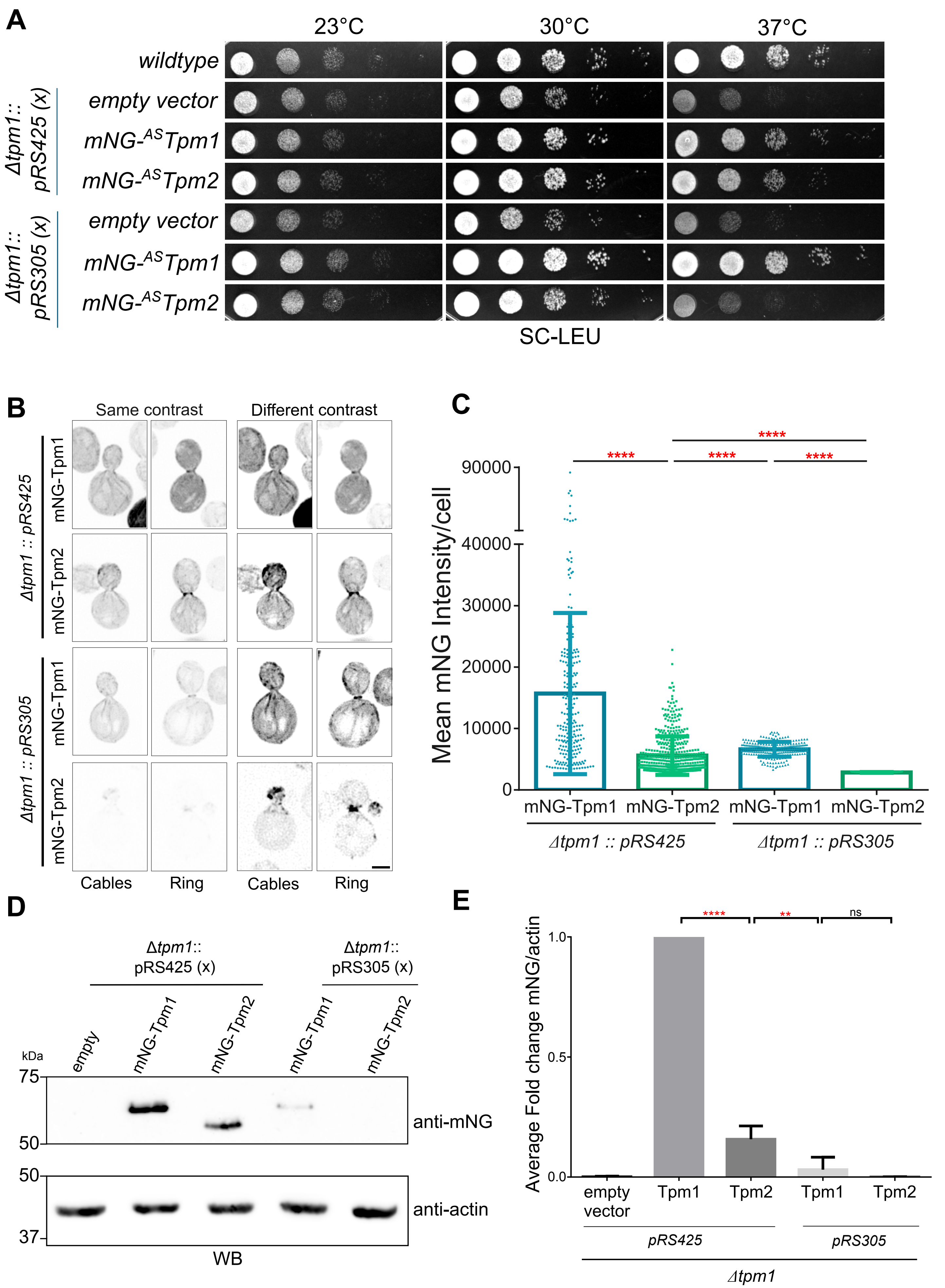
A
23°C
30°C
37°C
wildtype
empty vector
mNG-ASTpm1
mNG-ASTpm2
Δtpm1:: pRS425 (x)
Δtpm1:: pRS305 (x)
SC-LEU
B
Same contrast
Different contrast
mNG-Tpm1
mNG-Tpm2
Δtpm1 :: pRS425
Δtpm1 :: pRS305
Cables
Ring
C
Mean mNG Intensity/cell
90000
40000
30000
20000
10000
0
****
mNG-Tpm1
mNG-Tpm2
Δtpm1 :: pRS425
Δtpm1 :: pRS305
D
Δtpm1:: pRS425 (x)
Δtpm1:: pRS305 (x)
empty
mNG-Tpm1
mNG-Tpm2
kDa
75
50
37
anti-mNG
anti-actin
WB
E
Average Fold change mNG/actin
1.0
0.5
0.0
****
**
ns
empty vector
Tpm1
Tpm2
pRS425
pRS305
Δtpm1

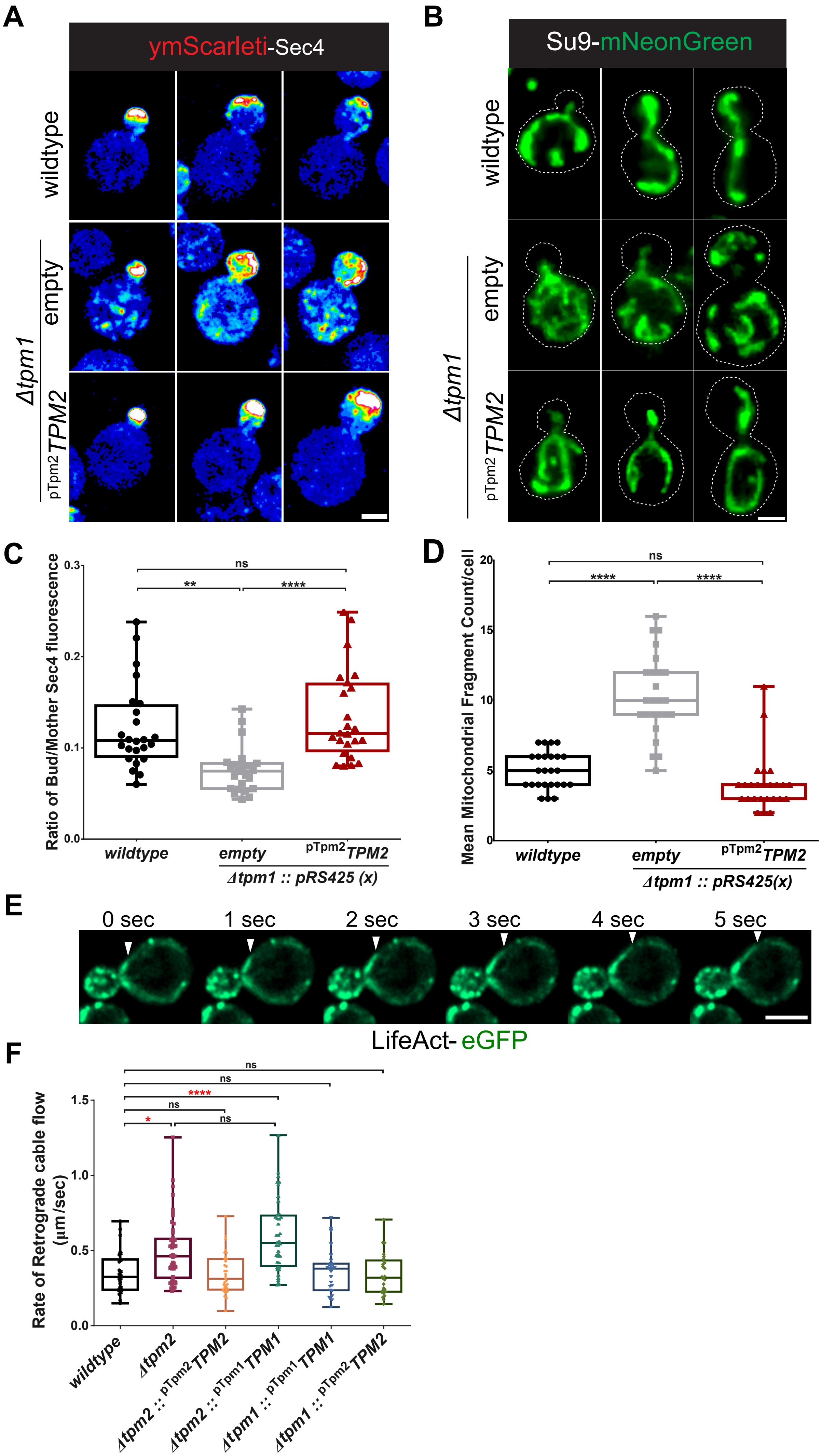
A
ymScarleti-Sec4
wildtype
Δtpm1
empty
pTpm2 TPM2
B
Su9-mNeonGreen
wildtype
Δtpm1
empty
pTpm2 TPM2
C
Ratio of Bud/Mother Sec4 fluorescence
0.0
0.1
0.2
0.3
ns
**
****
wildtype
empty
pTpm2 TPM2
Δtpm1 :: pRS425 (x)
D
Mean Mitochondrial Fragment Count/cell
0
5
10
15
20
ns
****
****
wildtype
empty
pTpm2 TPM2
Δtpm1 :: pRS425(x)
E
0 sec
1 sec
2 sec
3 sec
4 sec
5 sec
LifeAct- eGFP
F
Rate of Retrograde cable flow (µm/sec)
0.0
0.5
1.0
1.5
ns
ns
****
ns
*
ns
wildtype
Δtpm2
Δtpm2 :: pTpm2 TPM2
Δtpm2 :: pTpm1 TPM1
Δtpm1 :: pTpm1 TPM1
Δtpm1 :: pTpm2 TPM2

## A

Actin Cable quantification Workflow (adapted from *McInally et al., 2022 (bio-protocol)*)

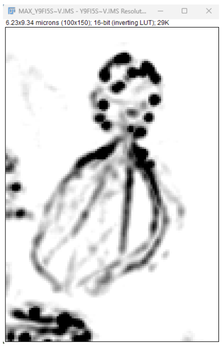

Inverted Grayscale image

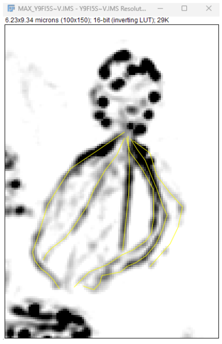

Traced actin cables in Fiji

Run script in Anaconda:

**actin_cable_length_analysis.py**

Adapted from McInally et al., 2022

## B

Mitochondrial Fragement Number quantification

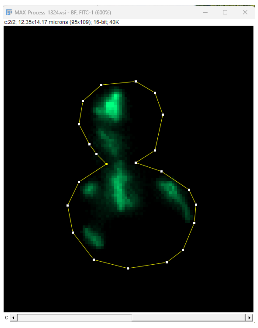

Mitochondrial signal

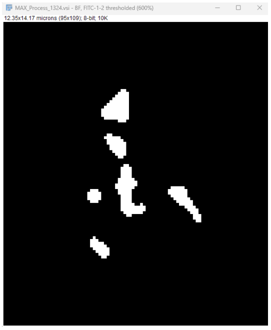

Segmented mitochondrial network using Mitochondria Analyzer Plugin

Measure mitochondrial fragment number in Fiji using "Analyze Particles"

2D-Threshold paramters used in Mitochondria Analyzer Plug-in

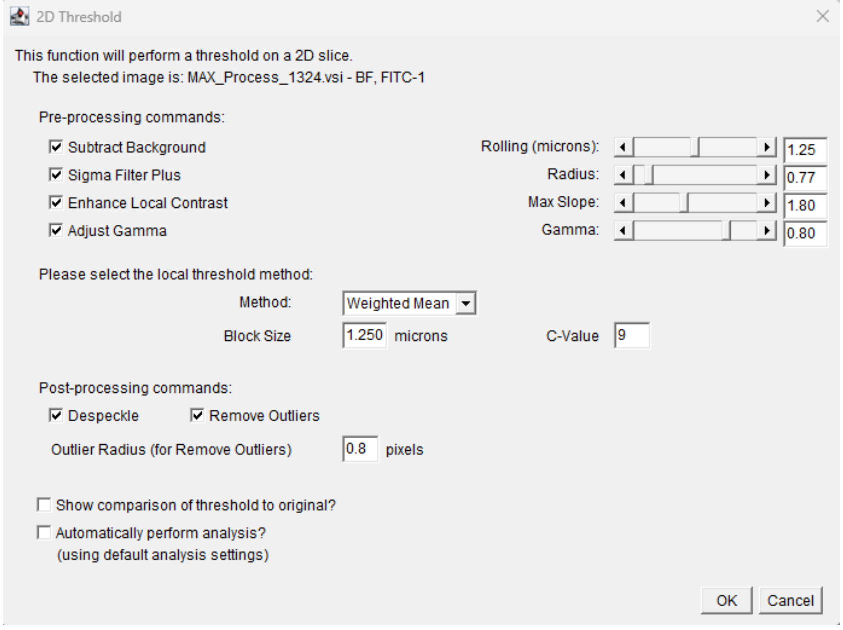

2D Threshold

This function will perform a threshold on a 2D slice.
The selected image is: MAX_Process_1324.vsi - BF, FITC-1

Pre-processing commands:

- [x] Subtract Background — Rolling (microns): 1.25
- [x] Sigma Filter Plus — Radius: 0.77
- [x] Enhance Local Contrast — Max Slope: 1.80
- [x] Adjust Gamma — Gamma: 0.80

Please select the local threshold method:

Method: Weighted Mean
Block Size 1.250 microns C-Value 9

Post-processing commands:

- [x] Despeckle
- [x] Remove Outliers

Outlier Radius (for Remove Outliers) 0.8 pixels

- [ ] Show comparison of threshold to original?
- [ ] Automatically perform analysis? (using default analysis settings)

OK Cancel
